# Engineering well-expressed, V2-immunofocusing HIV-1 envelope glycoprotein membrane trimers for use in heterologous prime-boost vaccine regimens

**DOI:** 10.1101/2021.07.20.453076

**Authors:** Emma T Crooks, Francisco Almanza, Alessio D’addabbo, Erika Duggan, Jinsong Zhang, Kshitij Wagh, Huihui Mou, Joel D Allen, Alyssa Thomas, Keiko Osawa, Bette T Korber, Yaroslav Tsybovsky, Evan Cale, John Nolan, Max Crispin, Laurent K Verkoczy, James M Binley

## Abstract

HIV-1 vaccine immunofocusing strategies have the potential to induce broadly reactive nAbs. Here, we engineered a panel of diverse, membrane-resident native HIV-1 trimers vulnerable to two broad targets of neutralizing antibodies (NAbs), the V2 apex and fusion peptide (FP). Selection criteria included i) high expression and ii) infectious function, so that trimer neutralization sensitivity can be profiled in pseudovirus assays. Initially, we boosted the expression of 17 candidate trimers by truncating gp41 and introducing a gp120-gp41 SOS disulfide to prevent gp120 shedding. “Repairs” were made to fill glycan holes and other strain-specific aberrations. A new neutralization assay allowed PV infection when our standard assay was insufficient. Trimers with exposed V3 loops, a target of non-neutralizing antibodies, were discarded. To try to increase V2-sensitivity, we removed clashing glycans and modified the V2 loop’s C-strand. Notably, a 167N mutation improved V2-sensitivity. Glycopeptide analysis of JR-FL trimers revealed near complete sequon occupation and that filling the N197 glycan hole was well-tolerated. In contrast, sequon optimization and inserting/removing other glycans in some cases had local and global “ripple” effects on glycan maturation and sequon occupation in the gp120 outer domain and gp41. V2 mAb CH01 selectively bound trimers with small high mannose glycans near the base of the V1 loop, thereby avoiding clashes. Knocking in a N49 glycan perturbs gp41 glycans via a distal glycan network effect, increasing FP NAb sensitivity - and sometimes improving expression. Finally, a biophysical analysis of VLPs revealed that i) ∼25% of particles bear Env spikes, ii) spontaneous particle budding is high and only increases 4-fold upon Gag transfection, and iii) Env+ particles express ∼30-40 spikes. Overall, we identified 7 diverse trimers with a range of sensitivities to two targets that should enable rigorous testing of immunofocusing vaccine concepts.

**Author Summary:** Despite almost 40 years of innovation, an HIV vaccine to induce antibodies that block virus infection remains elusive. Challenges include the unparalleled sequence diversity of HIV’s surface spikes and its dense sugar coat that limits antibody access. However, a growing number of monoclonal antibodies from HIV infected donors provide vaccine blueprints. To date, these kinds of antibodies have been difficult to induce by vaccination. However, two antibody targets, one at the spike apex and another at the side of the spikes are more forgiving in their ‘demands’ for unusual antibodies. Here, we made a diverse panel of HIV spikes vulnerable at these two sites for later use as vaccines to try to focus antibodies on these targets. Our selection criteria for these spikes were: i) that the spikes, when expressed on particles, are infectious, allowing us to appraise our vaccine designs in an ideal manner; ii) that spikes are easy to produce by cells in quantities sufficient for vaccine use. Ultimately, we selected 7 trimers that will allow us to explore concepts that could bring us closer to an HIV vaccine.

## Introduction

Contemporary HIV-1 vaccine candidates can routinely induce high titers of autologous tier 2 neutralizing antibodies (NAbs) (1–5). However, cross-neutralization is less predictable (6–8). This may be, in part, because vaccine NAbs generally target strain-specific gaps in envelope’s (Env’s) glycan armor, i.e. “glycan holes” (1, 5, 9, 10). By contrast, broad NAb (bNAb) targets are usually protected by neighboring glycans that nascent NAbs must evolve to avoid and/or bind (11–13).

Scores of bNAbs have been isolated from HIV-1-infected donors over the past decade, targeting 5 major conserved epitope clusters: the V2 apex, V3-glycan, CD4 binding site (CD4bs), gp120-gp41 interface/fusion peptide (FP) and the gp41 MPER (14). Exciting studies now show that NAbs with some breadth can be induced in vaccinated animals (7, 15, 16). Further efforts are needed to improve vaccine NAb breadth, titer and consistency.

Strategies to improve the quality of vaccine NAbs may be divided into 3 non-exclusive tracks (17–20). One is to trigger the expansion of bNAb precursor B cells (unmutated common ancestors: UCAs) (21–25). Vaccine-mediated triggering (26–32) may be improved by removing clashing glycans (3, 7, 33–35), reducing glycan size (13, 36–39), or by priming with core epitopes or scaffolds (8, 15, 16, 22, 32, 40–44). Ideally, priming creates an initial diverse pool of antibodies (Abs) that can be “filtered” by using carefully selected boosts to promote bNAb development (8, 34, 45). A second approach involves recapitulating natural bNAb development, using patient-derived Env clones to guide bNAb lineages to breadth (12, 24, 31, 46–53). This might be effective if UCAs are triggered by cognate Envs from the donors in which they arose (37, 38). A third strategy is to “immunofocus” NAbs, using trimers with high sensitivities to desired site(s) (3, 7, 8, 33–35, 40, 54).

Repeated immunization with the same Env trimer may cause NAbs to overly focus on distinct features of the vaccine strain, e.g., glycan holes. Trimer variants may be needed to drive NAb cross-reactivity. These modified trimers could be used in any of three regimen formats (2, 3, 10, 33–35, 54, 55). First, polyvalent mixtures of trimers could be used in each shot. This has been only modestly successful (54), perhaps because the most sensitive trimer amid the mixture dominates Ab responses. Moreover, while these mixtures are intended to promote cross-reactivity, they likely do not provide evolutionary direction for NAb development.

A second format is homologous prime-boosts, perhaps beginning with a highly sensitive trimer variant, followed by boosts with modified trimers of the *same* strain. This has been attempted with some success, typically by priming with trimers in which glycans surrounding the target are removed and boosting with trimers in which these glycans are reinstated (3, 7, 31, 34,35, 56, 57).

A third format, serial heterologous prime-boosts (SHPB), uses trimers from different strains in each shot (2, 54, 55, 58–60). This should eliminate strain-specific autologous NAbs. Success may hinge on whether nascent NAb-expressing memory B cells sufficiently cross-react with boosting trimers to keep them “on track”.

Here, we sought to assemble a panel of diverse trimers expressed on virus-like particles (VLPs) to simultaneously immunofocus both the V2 apex and fusion peptide (FP) epitopes. The resulting VLP SHPB regimens will later be tested in vaccine models, for example, the V2-specific CH01 “heavy chain only” (V_H_DJ_H_^+/+^) UCA knock in mouse (58). These two epitopes were chosen for several reasons. First, since the two sites do not overlap, the probability of inducing bNAbs is doubled. Second, both sites can accommodate multiple NAb binding modes (8, 42, 54) - this “plasticity” may increase the frequency of compatible germline Ab precursors. FP NAbs have common-in-repertoire features, and can be induced in many species, including mice (8, 15, 42). Some V2 NAbs (e.g. CH01, VRC38.01) also exhibit sufficiently common features, and thus do not depend heavily on rare V(D)J recombination and/or somatic hypermutation-driven events occurring during their formation, thus making them plausible vaccine prototypes (23, 25, 45).

V2 and FP NAbs both bind epitopes comprising of protein and glycan contacts. V2 bNAbs bind the N160 glycan and the neighboring basic C strand of a 5-strand β-barrel at the trimer apex (23, 61, 62) However, the binding may be regulated by protecting V1V2 glycans and long V1V2 loops (12, 33, 39, 54, 57, 63, 64). FP bNAbs, exemplified by VRC34, recognize the N-terminal gp41 fusion peptide, along with the proximal N88 glycan of gp120, but clash with gp41’s N611 glycan (8, 42, 43, 65, 66).

The preferential binding of CH01 and VRC34 to trimers produced in GnT1-cells, in which glycan maturation is blocked, suggests that both NAbs contact the stems of their respective glycans at positions N160 and N88, respectively, and that bulky glycan head groups hinder their binding. CH01 often mediates sub-saturating neutralization, even at high concentrations (13). This observation may stem from the differential glycosylation of otherwise identical trimers. Indeed, some sequons (glycosylation motifs) may be occupied by a variety of glycans or may be skipped altogether (67). This variability could have direct consequences on NAbs that either bind to trimers or are unable to bind due to glycan clashes. Glycan variation may also impact V1V2 folding (68). Since FP neutralization depends largely on the FP sequence, engineering trimers to maximize the induction of FP NAb breadth should be relatively straightforward. V2 bNAb ontogeny is more complex. In natural infection, a basic V2 C-strand (residues 166-171) promotes initial electrostatic NAb contacts. The C strand then becomes increasingly charge neutral as the virus escapes, promoting NAb reactivity with anchor residues, usually N160 glycan, and conserved residues at positions 168, 169, 171 and 173 of the C-strand, depending on the V2 NAb specificity (24, 36, 69).

We previously showed that VLPs expressing native JR-FL trimers, like their soluble, “near native” counterparts (e.g. SOSIP), regularly induce potent autologous NAbs (1, 10, 35). Advantages of SOSIP include its facile manufacture and rational, structure-driven vaccine improvements (70, 71). However, one drawback is that the V2 apex is less compact than on native trimers (72, 73), which may explain why they induce V2-specific Abs that differ markedly from V2 bNAbs (54, 70, 74). Second, the glycan-free base of SOSIP is an immunodominant, non-neutralizing target that could dampen responses to desired sites (75). Third, SOSIP partially exposes the V3 loop, leading to induction of V3 non-NAbs that are more modest against VLPs (1, 6, 35, 76–78). Fourth, SOSIP exhibits more unoccupied sequons and a more heterogeneous and immature glycoprofile compared to native trimers (9, 54, 67, 79–82), perhaps due to structural differences and/or to soluble trimer overexpression outpacing glycosylation machinery. Consequently, some SOSIP-induced Abs are unable to navigate membrane trimer glycans even on the cognate strain (9, 83, 84).

The transmembrane domain and cytoplasmic tail of membrane trimers anchor, stabilize, and modulate the external spike conformation, in particular the V2 apex (85, 86) in ways that cannot be readily achieved by soluble trimer. A further advantage of membrane trimer immunogens is that they can be directly checked for sensitivity in pseudovirus (PV) neutralization assays. This link to our desired endpoint (neutralization) eliminates any concerns that antigen binding assays are not anchored in functional relevance. Indeed, PV assays can reveal subtleties such as incomplete saturation mentioned above that might best be eliminated in candidate immunogens. Accordingly, we used NAbs CH01 (V2) and VRC34 (FP) as the main NAb probes to appraise candidate trimers, with particular emphasis on V2, due to its relative complexity.

Vaccine platforms worked well for their respective initial prototype strains, e.g. JR-FL for VLPs and BG505 for SOSIP. However, adapting these platforms to other strains is not straightforward. For SOSIP, many strains do not efficiently form well-folded near-native trimers, but mutations can remedy this problem, sometimes by leveraging BG505 sequences as “scaffolding” (87–91). In contrast, since membrane trimers are by definition native, proper folding is not usually a problem. However, expression is often insufficiently robust (92), but may be possible to improve (93–96). Accordingly, we took two approaches to generate our panel of trimers from 17 initial strains. First, we screened trimer expression of V2-sensitive strains. Second, we sought to increase V2 sensitivity of well-expressed strains. From there, we used a variety of repairs and adjustments to select a panel of 7 diverse, well-expressed VLPs for prospective SHPB vaccine studies.

## Results

Immunofocusing vaccine strategies entail priming with trimers exquisitely sensitive to desired NAbs, followed by boosting with trimers that are increasingly resistant to the same NAbs. Successive shots may reinstate clashing glycans absent in the prime and may concomitantly vary in the amino acid sequence of peripheral or variable parts of the epitope to encourage NAb focus on conserved “anchor” residues within the epitope(s). Successive use of trimer variants of the same strain may best mimic the conditions of bNAb development in natural infection (24, 36, 69). Conversely, changing strains in successive boosts should ensure a focus on common vulnerable sites, while minimizing the risk of strain-specific Abs. With these ideas in mind, we sought to identify a panel of trimers to immunofocus V2 and FP epitopes. Specifically, we sought well-expressed trimers with a range of CH01 sensitivities from at least 5 sequence-diverse strains to enable us to later test various prime-boost concepts in a mouse CH01 knockin model (58). Strains with broad sensitivity to multiple V2 NAbs are desirable (23). They should also be functional in pseudovirus (PV) assays, so that their neutralization sensitivity can be calibrated in a way that is directly relevant to our goal (inducing bNAbs). Finally, trimers should not be overtly sensitive to non-neutralizing V3 mAbs.

Given its relative complexity, we focused largely on identifying V2-sensitive trimers, with the intent to later “knock in” FP sensitivity, as needed. We previously assessed the trimer expression of various strains and found that there were vast expression differences (92). Poor expression is common - and is likely to be a problem for vaccine use. Accordingly, we took two approaches to identify well-expressed V2-sensitive trimers. First, we evaluated several known V2-sensitive strains, based on compiled neutralization data (97) (group 1), reasoning that some may be naturally well-expressed and that we may be able to improve the expression of others, as needed. In a second, complementary approach, we sought to engineer well-expressed membrane trimers to knock in V2 sensitivity (group 2). Key features of 17 candidate strains, including 12 group 1 strains and 5 group 2 strains are shown in Fig. 1. Aligned and annotated sequences are shown in S1 Fig.

**Figure 1.**
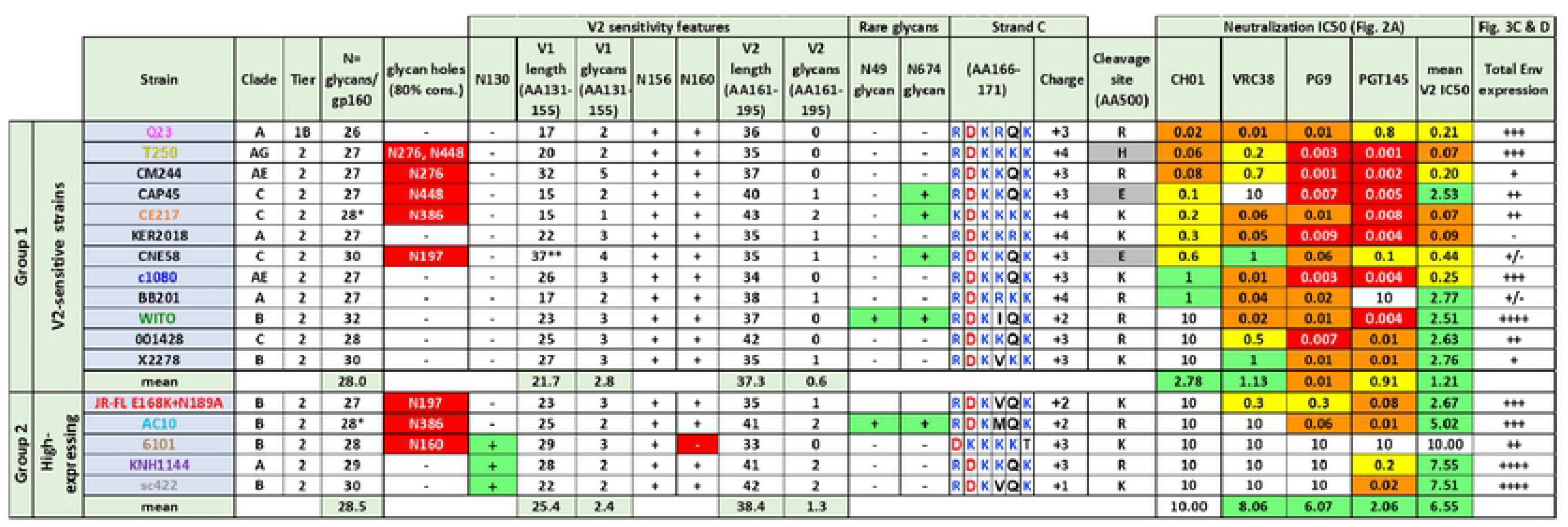
Key features of candidate Env strains. 17 strains were placed in two groups: i) 12 group 1 strains are naturally sensitive to multiple V2 NAbs; ii) 5 group 2 strains exhibit high membrane trimer expression. Strain names are abbreviated (see Methods). An asterisk in total glycans per gp160 protomer indicates overlapping sequons in CE217 (N396 and N398) and AC10 (N190 and N191), only one of which can carry a glycan. Glycan holes are listed whenever a ≥80% conserved glycan is absent. V2 sensitivity features are shown, including glycans involved in nAb binding or clashes, loop lengths. A double asterisk for the CNE58 V1 loop denotes a possible internal hairpin disulfide loop (Fig. S1). Rare glycans N49 and N674 are shown. Strand C sequence (AA166-171) is shown with basic residues in blue and acidic residues in red, along with overall strand C charge. The amino acid at position 500 may influence gp120/gp41 processing (gray highlights non-lysine or arginine residues). PV IC50s for V2 NAbs (CF2 assay) (see Fig. 2A). JR-FL neutralization data is for the E168K+N189A mutant. Total Env expression, as judged by SDS-PAGE-Western blot (see Fig. 3C and D).

Group 1 strains all carry glycans at positions N156 and N160, a K/R-rich basic C strand (residues 166-171) and lack the clashing N130 glycan at the V1 loop base (S1 Fig). Several group 1 strains, including Q23, WITO and c1080 have few other V1V2 glycans aside from the apex N156 and N160 glycans and have short V1 lengths - features that may be hallmarks of high V2 sensitivity (21, 22, 54, 57). Group 2 includes five strains that previously expressed well as gp41 cytoplasmic tail-truncated (gp160ΔCT) trimers on VLP surfaces (Fig. 1 in (92)). Group 1 trimers show higher sensitivity to 4 prototype V2 bNAbs (Fig. 1), of which Q23, T250, c1080 and WITO were of exceptional interest, in being both V2-sensitive and well-expressed (Fig. 1). Previously, E168K+N189A mutations ‘knocked’ V2 sensitivity into the otherwise resistant JR-FL strain (13, 98). This pair of mutations introduce a C strand lysine and eliminate a competitive glycan in the distal part of the V2 loop (V2’; AA 180-191; S1 Fig). Unlike the JR-FL parent, this mutant is sensitive to 3 out of 4 prototype V2 NAbs (Fig. 1), demonstrating the feasibility of modifying well-expressed group 2 trimers to knock in V2 sensitivity.

The number of glycans per gp160 protomer varies between strains (Fig. 1). Counting non-overlapping sequons in an HIV-1 Env alignment of 4,582 diverse sequences in the LANL database between amino acids 31-674 (99), the median sequons/protomer is 29 (inter-quartile range of 28-31). Glycans have two major functions. Structural glycans e.g. N262 (100) impact folding and are therefore relatively conserved (shaded blue In S1 Fig. When these glycans are missing, Ab-sensitive “glycan holes” are typically created. In keeping with the role of glycans in folding, it is of interest that well-expressed group 2 trimers have an average of 1 more glycan per gp160 protomer than group 1 trimers (Fig. 1). In contrast to conserved structural glycans, other glycans are commonly gained or lost to facilitate NAb escape ((101); shaded yellow in S1 Fig).

Several of the 17 strains exhibit “glycan holes” (Fig. 1, orange boxes in S1 Fig). Filling these holes with glycan may eliminate vulnerable strain-specific sites that may delay NAbs developing to the intended target(s) (1, 9, 10, 83, 99). If a threshold number of glycans are needed for high expression, then filling glycan holes may compensate for removing clashing glycans at targeted sites. Two strains (WITO and AC10) exhibit rare glycans at positions 49 and 674 that could be related to their high expression (Fig. 1). One final variable that may be relevant to our goals is that some strains lack a basic residue at position 500 near to the furin processing site (Fig. 1), which could adversely impact gp120/gp41 maturation. Below, we explore the effects of modifying these and other Env features to develop a panel of well-expressed immunofocusing membrane trimers for vaccine use.

### V2 sensitivity of trimer panel candidates

We first measured sensitivities to 4 prototype V2 NAbs (Fig. 2A), along with the CH01 Unmutated Common Ancestor (UCA) and a germline-reverted form of VRC38 (both termed ‘UCA’ hereafter, for brevity) (23). V3 non-NAb sensitivity was used to monitor trimer folding, so that loosely folded trimers that overtly expose V3 epitopes can be discarded. Although V3 mAbs 14e and 39F did not neutralize any of our strains, it is important to verify that their epitopes are actually present should the trimers be ‘triggered’ upon CD4 binding (13). To do this, we mixed PVs with soluble CD4 (sCD4) and measured the ability of V3 mAbs (39F and 14e) to inhibit infection of CCR5-expressing cells (102). In this format, V3 mAbs neutralized 11 of our strains (S2 Fig). CE217 was later verified as 14e-sensitive in mutation analysis (see below). 14e-sensitivity of the other 5 strains of our panel could not be confirmed, as their infectivities were too low.

**Figure 2.**
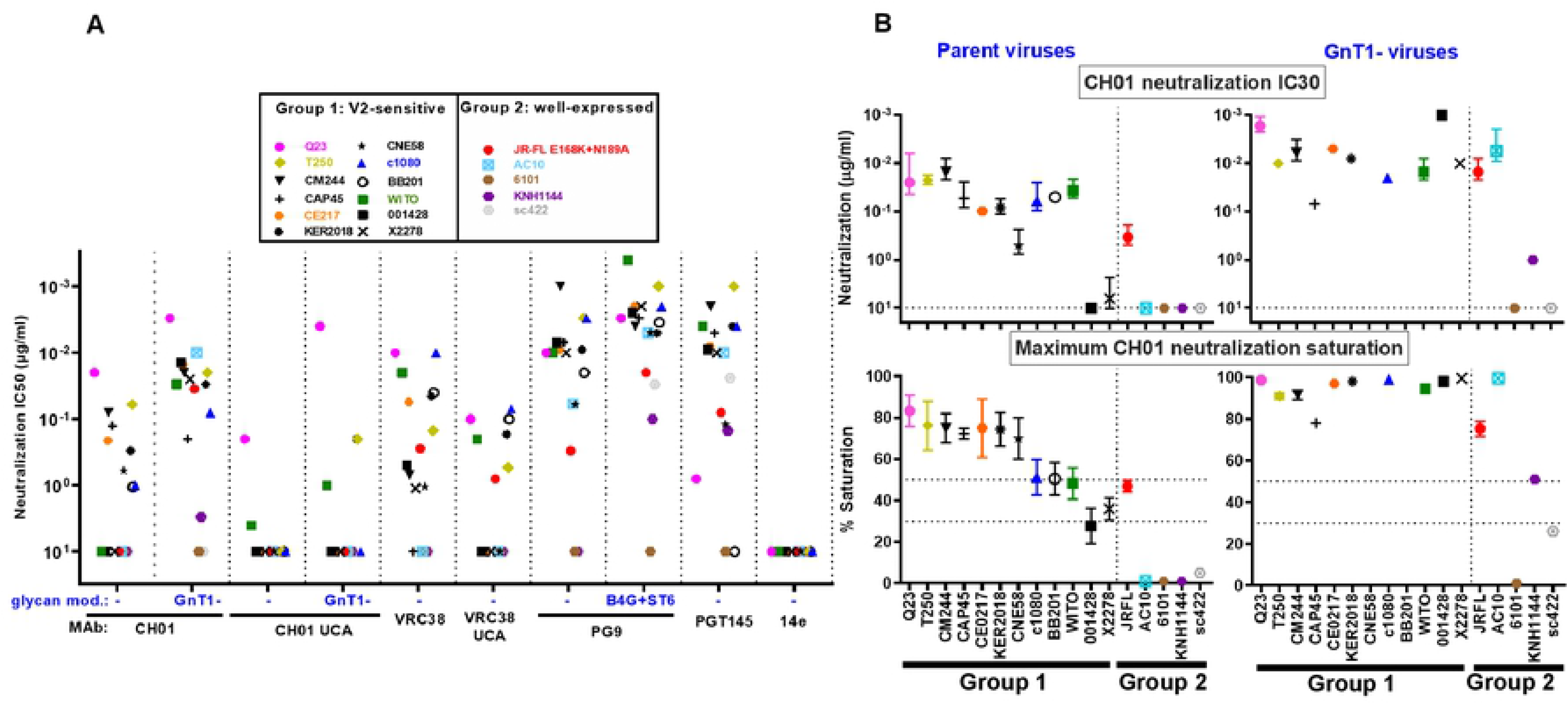
V2 NAb sensitivity of candidate strains. A) MAb IC50s against 17 candidate PVs bearing full-length wild-type (WT) gp160 spikes, except for WITO, AC10, 6101, KNH1144 and sc422, that were gp41 cytoplasmic tail-truncated (gp160ΔCT). For CH01 and VRC38, UCA sensitivities are shown. CH01 and PG9 NAb IC50s were measured against PVs bearing Envs with engineered glycans: GnT1- and B4GalT1+ST6Gal1 (abbreviated as B4G+ST6), respectively. GnT1-PV data BB201 and CNE58 strains are not shown, as infection was insufficient. B) CH01 IC30s and % maximum CH01 neutralization saturation in unmodified (left) and GnT1-(right) formats.

For CH01 and its UCA, we also measured IC50s against PV produced in GnT1-cells. We previously showed that CH01 saturation improves against GnT1-PV, presumably as clashes with larger glycan head groups are eliminated (13). Similarly, PG9 neutralization is more effective against B4GalT1+ST6Gal1-(abbreviated “B4G+ST6”) modified PV that increases hybrid glycans and terminal α-2,6 glycan sialylation (13). Neither of these modifications overtly increase V3 sensitivity, suggesting that trimer folding is not impacted.

Q23 was the most sensitive strain to MAbs CH01, VRC38 and their UCAs (Fig. 2A; (23)) and was also highly PG9-sensitive. Although Q23’s tier 1B classification and moderate PGT145-sensitivity reflect a somewhat less compact V2 apex compared to other strains (Fig. 1), it is nevertheless 14e-resistant, and may therefore be useful to prime V2 NAbs in a vaccine regimen.

Surprisingly, WITO, 001428, and X2278 were not neutralized by CH01 10μg/ml (Fig. 2A), contrasting previous data (97). This may be due to CH01’s characteristic “sub-saturating” neutralization and/or that our ‘workhorse’ CF2 assay is slightly less sensitive than the commonly used TZM-bl assay (13). Indeed, CH01 exhibited IC30s of 0.04μg/ml and 6μg/ml, respectively against WITO and X2278 (Fig. 2B, top left) and almost reached an IC30 against 001428 at 10μg/ml (Fig. 2B, bottom left).

In contrast, 13 of 15 GnT1-PVs were CH01-sensitive (Fig. 2B, top right; BB201 and CNE58 GnT1-PVs did not infect sufficiently). For 10 of the CH01-sensitive GnT1-PVs, maximum CH01 saturation was close to 100% (Fig. 2B, bottom right) and its IC30s were also lower (Fig. 2B, compare upper panels). VRC38, PG9 and PGT145 neutralized all of the group 1 strains, except that PGT145 did not neutralize BB201. In several cases, they also neutralized group 2 strains (Fig. 2A). Only 6101 was resistant to all V2 NAbs, probably due to its missing N160 glycan (Fig. 1, S1 Fig).

Overall, Q23’s exquisite sensitivity to CH01, its UCA and other V2 NAbs support its use as a V2 priming immunogen. Several other CH01-sensitive strains could be useful in boosting. However, as outlined below, selected strains may benefit from engineering to increase V2 sensitivity and/or trimer expression.

### *In situ* membrane expression of candidate trimers

Trimer expression on VLP surfaces is improved by truncating gp160 at position 708, leaving a 3 amino acid gp41 tail (gp160ΔCT; Fig. S1, Fig. 3A (compare lanes 1 and 2)). Based on our previous work, high VLP trimer expression is achieved by co-transfecting Env plasmids with MuLV Gag (Fig. S1 in (23)) and Rev plasmids (used when Env is not codon-optimized). The SOS mutation (501C+605C) further improves JR-FL trimer expression (Fig. 3A, compare lanes 2 and 3). E168K and E168K+N189A variants of JR-FL gp160ΔCT SOS were also well-expressed (Fig. 3A, lanes 4 and 5). Gp160ΔCT SOS mutants of strains Q23.17 and CNE58 were also better expressed than their gp160 WT and gp160ΔCT WT counterparts (Fig. 3A, lanes 6-10), although neither expressed as well as JR-FL. One explanation for the high SOS trimer expression is that gp120 shedding is eliminated, evidenced by the lack of gp41 stumps (Fig. 3A, compare lanes 2 and 3).

**Figure 3.**
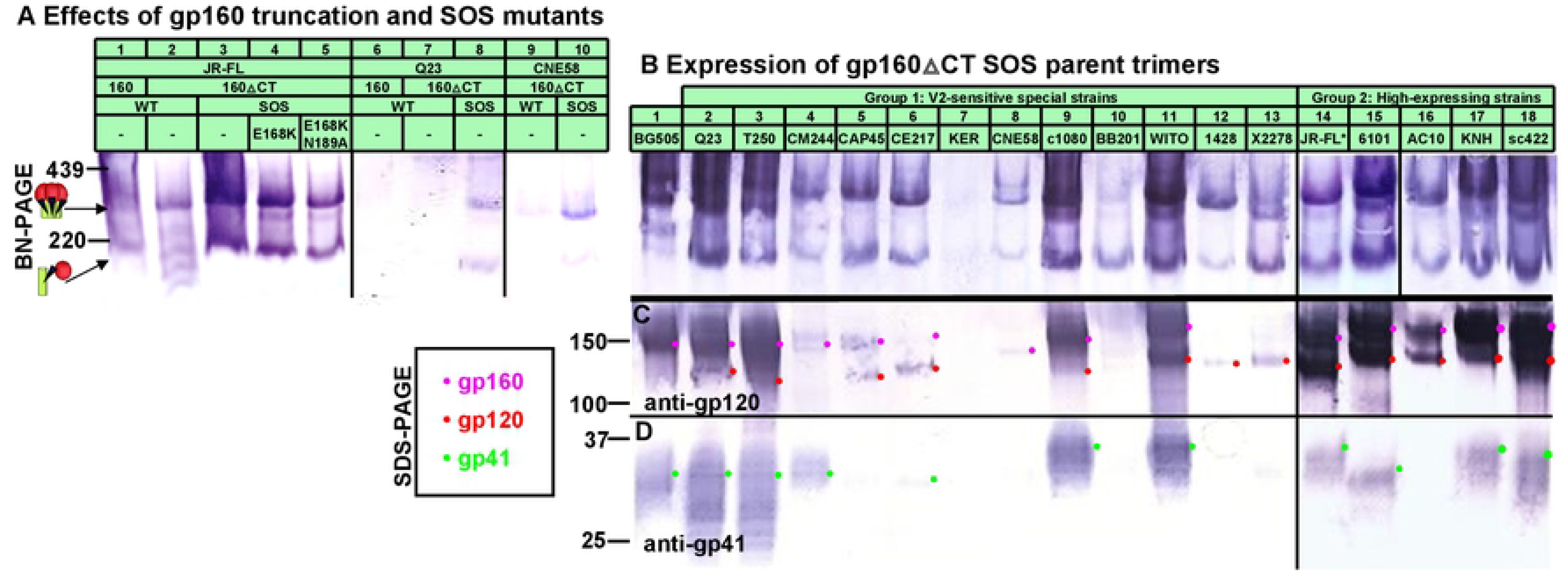
Gp160ΔCT and SOS mutations improve expression of candidate strains. A) VLP trimer expression with or without gp41 truncation (gp160ΔCT) and SOS mutations, probed with anti-gp120 and anti-gp41 mAb cocktail. SOS gp160ΔCT trimer expression of candidate strains visualized by B) BN-PAGE-Western blot and by SDS-PAGE-Western blot, probing with anti-gp120 (C) or anti-gp41 (D) mAb cocktails. All Envs were expressed using robust expression plasmids (pVRC8400 or pCDNA3.1), except for Q23 in part A lanes 6-8 and BB201 in part B lane 10, for which pCR3.1 was used. A high expression plasmid (pVRC8400) was used to express Q23 in part B, lane 2.

SOS and gp160ΔCT mutations were made in Env plasmids previously used to make PV of the 17 candidate strains (Fig. 1), along with BG505 as a reference. SOS gp160ΔCT Env expression was examined in BN-PAGE- and SDS-PAGE-Western blots (Fig. 3B-D). BG505 trimers expressed modestly well (Fig. 3B, lane 1) and consisted mainly of uncleaved gp160 (Fig. 3C, lane 1). Although gp120 was not visible (Fig. 3C, lane 1), some gp41 was detected (Fig. 3D, lane 1).

Q23 SOS gp160ΔCT expression by the pVRC8400 plasmid (Fig 3B-D, lane 2) was far higher than by the pCR3.1 plasmid (Fig. 3A, lane 8). Poor BB201 Env expression may also be due to use of the pCR3.1 plasmid (Fig. 3B-D, lane 9). For all other strains, expression plasmids pVRC8400 or pCDNA3.1 were used. Group 1 strains T250, c1080, and WITO expressed high levels of trimer, gp160, gp120 and gp41 (Fig. 3B-D, lanes 3, 9 and 11). Strains CM244, CAP45, CE217, CNE58, 001428 and X2278 expressed modestly (Fig. 3B-D, lanes 4, 5, 6, 8, 12 and 13). The remaining group 1 strain, KER2018, expressed extremely poorly (Fig. 3B-D, lane 7). We used SDS-PAGE-Western blots to monitor total Env output from hereon.

In contrast to the mixed expression of group 1 strains, all 5 group 2 SOS gp160ΔCT mutants had high levels of trimer (Fig. 3B, lanes 14-18). Gp160 and gp120 expression was high in all cases (magenta and red dots in Fig. 3C). Corresponding gp41 bands were also observed in all but AC10. Overall, these blots reveal vastly different Env expression of different strains that could greatly impact their utility in vaccines (92). The group 2 strains and the 4 group 1 strains that express high levels of functional authentic gp120/gp41 trimers (Q23, T250, c1080 and WITO) are of particular interest for follow up. Below, we used various strategies to modify the most promising strains to knock in V2 sensitivity and/or to improve Env expression.

#### JR-FL modifications

We first modified our prototype vaccine strain, JR-FL, to try to improve its V2 and FP sensitivity. The E168K+N189A mutant was sensitive to VRC38, PG9 and PGT145 (Fig. 1) and partially CH01-sensitive (Fig. 2B). V2 sensitivity might be improved by removing V1V2 clashing glycans and/or by increasing strand C’s overall basic charge. To maintain high expression, modifications were made in the JR-FL SOS gp160ΔCT background. We initially compared WT and SOS PV NAb sensitivities. As shown previously, SOS PV infection can proceed after receptor engagement by adding a low molarity reducing agent to break the gp120-gp41 disulfide (103). WT and SOS PV neutralization profiles were broadly similar (S3A Fig). However, CH01 saturation of SOS mutant PV was greater and an IC50 was measurable. Thus, the SOS mutant improves trimer expression, retains V3 resistance and slightly improves CH01 sensitivity.

Since the JR-FL N188 and N189 sequons overlap, only one site can be occupied by glycan. We therefore investigated the effects of knocking out each sequon alone or together. Despite only moderate effects on V2 IC50s (Fig. 4, lanes 1-3), both single mutants improved PG9 saturation. However, the double mutant exhibited reduced PG9 sensitivity and saturation (S3B Fig, Fig. 4B lane 19). Unlike the E168K, E168K+N188A and E168K+N188A+N189A mutants, the E168K+N189A mutant was modestly CH01-sensitive (Fig. S3A, Fig. 4B, lanes 1-3, 19). We therefore used JR-FL E168K+N189A as a “parent” clone to overlay further mutations, then monitor Env expression (monitored by SDS-PAGE-Western blot), PV infectivity (Fig. 4A), and NAb sensitivity (Fig. 4B).

**Figure 4.**
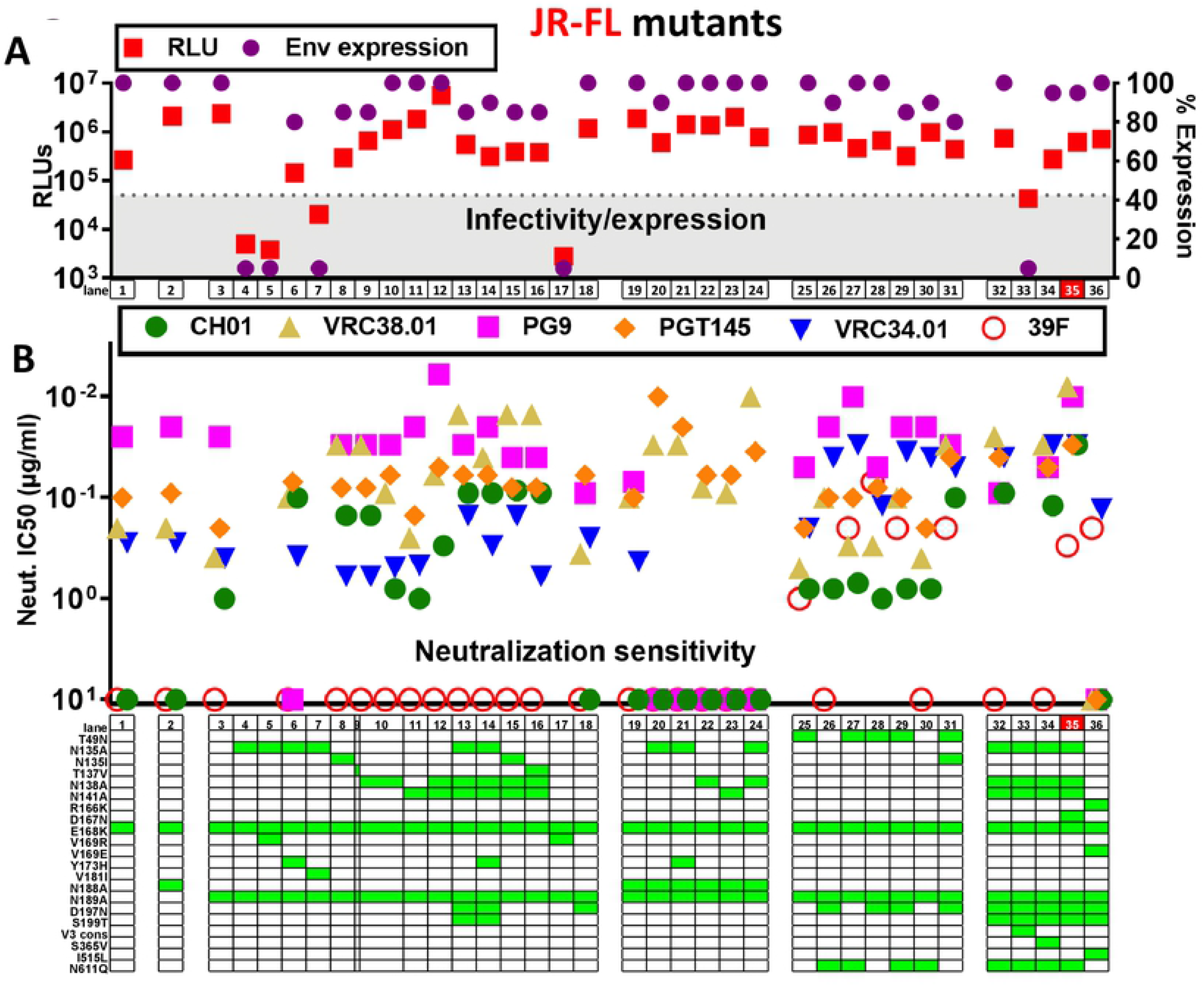
Effect of mutations on JR-FL SOS gp160ΔCT trimer expression, infectivity and NAb sensitivity. Effect of mutations on A) JR-FL gp160ΔCT SOS trimer infectivity, and total Env expression (quantified by SDS-PAGE-Western blot), and B) mAb sensitivity. The mutant with the most desirable features is highlighted in red (lane 35). The V3 consensus (V3 cons) mutant in lane 33 included H310R+R315Q+T319A+E322D mutations.

V2 NAb sensitivity typically depends on glycans at positions N156 and N160 and basic (K/R) residues in strand C (54, 57, 61, 62, 98) (S4 Fig). K168 and K171 are relatively conserved V2 NAb “anchor” residues (23, 61); K/R residues at 166, 169 and 170 also contribute to strand C’s net positive charge (S4 Fig), boosting electrostatic interactions with V2 NAbs. Our JR-FL E168K+N189A mutant contains these anchor residues and a net charge of +2, which is somewhat lower than many V2-sensitive strains that, unlike JR-FL, exhibit K/R at position 169 of strand C (Fig. 1). We return to this point later below.

### V1 and V2 loop glycan deletions

We initially sought to maximize JR-FL’s CH01 sensitivity, hoping to concomitantly boost sensitivity to other V2 NAbs. We first removed potentially clashing glycans at positions 135, 138 and 141 at the V1 loop base (S1 Fig). The N135A mutant dramatically reduced PV infectivity and trimer expression (Fig. 4A, lane 4), suggesting a folding problem. To try to obtain an infectious N135A mutant, we combined it individually with 3 local V2 substitutions commonly found in other strains (S4 Fig). A V169R mutation would increase strand C positive charge, possibly improving V2 sensitivity. However, neither the N135A+V169R double mutant nor the V169R alone were infectious or expressed trimer (Fig. 4A, lanes 5 and 17). Y173 and Y177 of the V2 loop (S4 Fig) may interact with residues N300 and K305 at the V3 loop base to influence trimer folding (104). H173 is also common (S4 Fig). N135A+Y173H restored some PV infectivity, albeit not to the level of the parent (Fig. 4A, lane 6). Moreover, it dramatically improved CH01 sensitivity and saturation (Fig. 4B, lane 6, S5A Fig). Conversely, PG9 sensitivity was eliminated (Fig. 4B, lane 6). Since the Y173H mutant alone did not affect CH01 sensitivity (S5A Fig), we infer that the increased CH01 sensitivity of N135A+Y173H is due to N135A. N135A+V181I also improved infectivity, albeit insufficiently to measure PV sensitivity (Fig. 4A, compare lanes 4 and 7). In an alternative approach, we made mutants N135I and T137V. These both improved CH01 and VRC38 sensitivity, but unlike N135A+Y173H, PG9 sensitivity was retained (Fig. 4B, compare lanes 3, 8 and 9). Compared to N135A+Y173H, these mutants were slightly more VRC38-sensitive, but slightly less CH01-sensitive (Fig. 4B compare lanes 6, 8 and 9). Overall, N135I and T137V were preferable to N135A+Y173H as they are better expressed and more infectious.

N138A did not impact trimer expression or infectivity (Fig. 4A, compare lanes 3 and 10) and modestly improved CH01 sensitivity and saturation, but not to the extent of N135 glycan knock out mutants (Fig. 4B, compare lanes 3, 6, 8, 9 and 10; S5A Fig). VRC38 and PGT145 sensitivity was also higher, but PG9 sensitivity was unchanged (Fig. 4B, lanes 3 and 10). The greater impact of N135A compared to N138A on V2 NAb sensitivity is consistent with its closer proximity to the core N156 and N160 glycans (Fig. 5A).

**Figure 5.**
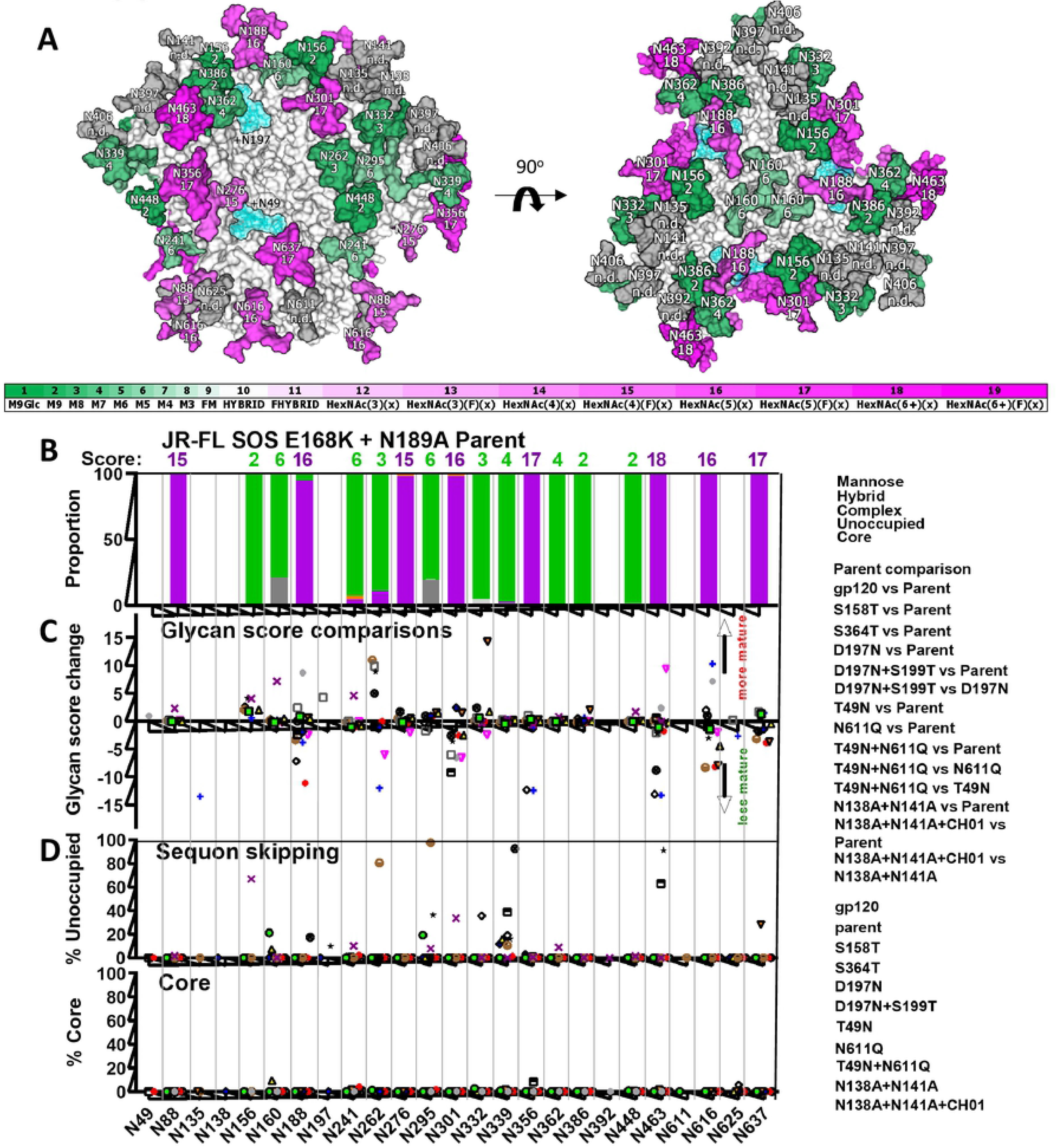
Effects of mutants on JR-FL membrane trimer glycan maturation and occupation. Related to S1 Data and Analysis, S7 Fig and S8 Fig. A) In a trimer model (pdb: 6MYY), each glycan is numbered according to the prototype HXB2 strain (see S1 Fig) and is given a maturation score, derived from LC-MS analysis of parent JR-FL SOS E168K+N189A VLPs (S1 Data and analysis). Glycans are colored in shades of green (high mannose) or magenta (complex), according to their score. Thus, untrimmed high mannose glycans are dark green and trimmed high mannose glycans are shown in lighter hues of green. Conversely, heavy complex glycans are shown in dark magenta, whereas smaller complex glycans are shown in lighter hues of magenta. Some glycans, rendered in gray, were not resolved in the JR-FL parent and therefore have no glycan score (not done; n.d.). Glycans at positions N49 and N197 are modeled as blue translucent masses. B) Glycan identity and scores at each sequon in JR-FL SOS E168K+N189A VLPs determined by LC-MS. Glycans were assigned scores by their degree of maturation (S7 Fig). C) Changes in glycan scores at each position between sample pairs. A negative score implies a shift to less mature glycan and *vice versa*. Data are only shown at positions where a glycan was detected in >10% of the peptides from both samples. Calculations for the score differences are shown in S1 Data and analysis and are modeled in S8 Fig. D) Sequon skipping and core glycans at each position.

N141A had little effect on any parameter (Fig. 4, compare lanes 3 and 11). However, if the N141 sequon of JR-FL membrane trimers is only 50% occupied by glycan, as reported previously (79), the negligible effect of N141A is perhaps not surprising. Incomplete N141 occupation may be due to spatial competition with the N138 sequon. If so, N141 occupation may increase when N138 is absent. Accordingly, a N138A+N141A double mutant exhibited improved CH01, VRC38 and PG9 sensitivity compared to N138A alone (Fig. 4B, lanes 10-12, Fig. S5A). Infectivity and trimer expression were also comparable to that of the parent (Fig. 4A, lanes 3 and 12).

To try to further improve CH01 sensitivity, we removed all 3 V1 glycans together, first as a N135A+N138A+N141A mutant. To compensate for the loss of 3 glycans and to also fill in a well-known JR-FL glycan hole, we overlaid a D197N+S199T mutation (1). Located at the base of the V2 loop, this glycan could impact V2 NAb binding (Fig. 5A). The N135A+N138A+N141A+D197N+S199T mutant was well-expressed and infectious. Unlike the single N135A mutant, an additional Y173H mutation was not needed (Fig. 4A, lanes 13 and 14). Thus, either the absence of other V1 glycans and/or the added N197 glycan compensated for the folding defect of the N135A mutant (Fig. 4A, lane 4). The triple V1 glycan mutant was marginally more sensitive than N135 single glycan knockout mutants (Fig. 4B, compare lanes 6, 8, 9 to lanes 13 and 14; S5A Fig). Since the N197 glycan lies at the edge of the V2 apex (Fig. 5A), it is possible that the D197N+S199T mutant used in combination with the triple V1 glycan mutant may directly impact V2 NAb sensitivity. Compared to the parent, the D197N mutant alone exhibited higher PGT145 sensitivity but weaker or no PG9 and CH01 sensitivity, respectively (Fig. 4B, lanes 3 and 18). This suggests that the D197N glycan knock in alone does not account for the high V2 sensitivity of the N135A+N138A+N141A+D197N+S199T mutant. Triple V1 glycan knockout mutants using N135I and T137V were also highly V2-sensitive, (Fig. 4B, lanes 13-16). These latter mutants do not contain a D197N mutation, further suggesting that most gains in V2 sensitivity ascribe to V1 glycan deletion, not N197 glycan addition.

We next tested if some of the above mutants might be further augmented by removing *both* the N188 and N189 glycans (Fig. 4, lanes 19-24). In this context, N135A showed high VRC38 and PGT145 sensitivity (Fig. 4B, compare lanes 6 and 21). However, these mutants were neither CH01-nor PG9-sensitive. Therefore, the reduced PG9 saturation of the E168K+N188A+N189A mutant noted above (S3B Fig), was exacerbated by V1 glycan removal, suggesting that the N188 glycan is required for broad V2 NAb sensitivity V2.

### Improving FP nAb sensitivity

We next engineered changes to improve FP NAb sensitivity. Specifically, we removed the N611 glycan clash with VRC34 (8). We also knocked in the rare N49 glycan (modeled in Fig. 5A) that is carried by well-expressed strains WITO and AC10 (S1 Fig), hoping to boost JR-FL trimer expression. Our model places the N49 glycan in between the N276 and N637 glycan sites (Fig. 5A). Its proximity to N637 led us to wonder if it might impact the local glycan network that regulates sensitivity to FP NAbs like VRC34 that contact N88 and clash with N611 glycans. We therefore toggled the N49 and N611 glycans, with or without D197N. T49N and N611Q glycan mutants did not appreciably impact PV infectivity or trimer expression (Fig. 4A, lanes 25-31). N611Q improved VRC34 sensitivity, while T49N did so to a lesser extent (S5B Fig). T49N+N611Q was only marginally more VRC34-sensitive than N611Q alone (Fig. 4B, lanes 3, 25, 27 and 30). T49N mutants were all modestly 39F-sensitive (Fig. 4B, lanes 25, 27-29, 31, S5C Fig), but remained PGT145- and PG9-sensitive. An additional N135I mutation concomitantly increased VRC38 and CH01 sensitivity (Fig. 4B, compare lanes 28 and 31). Overall, despite causing partial V3 exposure, the N49 glycan knock in did not perturb V2 apex epitope integrity and modestly improved VRC34 sensitivity.

### Effects of mutations on glycan maturation and occupation

S6A Fig shows an SDS-PAGE-Western blot of some of the above mutants to try to better understand the basis of the effects observed above (Fig. 4) and in reference to a structural model (Fig. 5A). Notably, N49 mutants caused a slight gp41 mass decrease, coupled with the expected slight gp120 mass increase (S6A Fig, compare lanes 1 and 2). The gp41 mass decrease of the T49N mutant was smaller than that of the N611Q mutant that knocks out a gp41 glycan (S6A Fig, lanes 1, 3, 5 and 6). As expected, the D197N mutant increased gp120 mass, but unlike T49N, did not impact gp41 mobility (S6A Fig, lanes 2 and 4). T49N+D197N led to a bigger gp120 mass increase than either of the single mutants, suggesting that glycans were added at both sites (S6A Fig, lanes 2, 4-6). Conversely, T49N+N611Q did not reduce gp41 mass further than N611Q alone (S6A Fig, lanes 5 and 6). Thus, the effect of T49N on gp41 is eliminated when combined with N611Q. V1 glycan knockout mutants revealed decreases in gp120 mass that were consistent with removing one or all 3 glycans (S6A Fig, lanes 7-10). However, gp41 mass did not change, as expected. Expression of these V1 mutants was somewhat weaker than the parent, most evident from the decreased gp41 staining (S6A Fig, lanes 7-10, Fig. 4A).

We next tested the impact of the N49 glycan knock in on gp41 sensitivity to endoglycosidase H (endo H). As reported previously (13), parent JR-FL gp41 was endo H-resistant, consistent with complex, fucosylated glycans. However, N49 mutant gp41 exhibited a ladder of endo H-sensitive species, consistent with the idea that the N49 glycan limits gp41 glycan maturation (S6B Fig, lanes 2 and 4). While the N49 glycan is close to gp41 glycan N637 (Fig. 5A), it is not close to the N611 glycan that clashes with VRC34. In one scenario, reduced N637 glycan maturation might allow greater N611 glycan flexibility, improving VRC34 binding. We further investigate the effect of the N49 glycan below.

To gain further insights into the effects of the various mutations above that add remove or modify particular sequons, we assessed glycan occupation and maturation by glycopeptide in-line liquid chromatography mass spectrometry (LC-MS) (67, 81). Each glycan type was given a score from 1 to 19, depending on the average maturation state (S1 Data and analysis, S7 Fig). Thus, the untrimmed high mannose glycan, M9Glc, has a score of 1, while the most highly branched and fucosylated complex glycan HexNAc(6+)(F)(x) has a score of 19. Glycan maturation scores of parental E168K+N189A trimers are modeled in Fig. 5A. Glycan scores and diversity at each site are summarized in Fig. 5B. The nature of glycans at each site generally match a previous report that categorized JR-FL PV Env glycans by another method (79). However, the N160 and N386 glycans were mostly high mannose in our hands but were complex by the other method.

We next evaluated glycan score differences at each site in pairs of samples. Score changes were recorded in a dot plot (Fig. 5C) for sites that were >10% occupied by glycans (excluding core glycans) in both samples. Score differences for each pair are modeled (S8 Fig). Sequon skipping and core glycans are shown in Fig. 5D.

We first compared two preparations of JR-FL SOS E168K+N189A VLP trimers (’parent’) analyzed on different days to gauge sample and assay variation (S1 Data and analysis, S7 Fig, S8 Fig). This revealed minor differences in gp120 glycans, with a modest difference high mannose trimming at position N156 (Fig. 5C, S8 Fig). Gp41 glycans were all heavy and complex (S1 Data and analysis, S7 Fig). Sequon skipping was rare and varied between samples, occurring at positions N156 (0.87%) and N362 (0.4%) in one sample and at N160 (21.24%) in the other (S1 Data and analysis, average % skipping shown in Fig. 5D). Glycan core was found occasionally (∼5% or less) at 4 sites in one sample, but not at all in the other (S1 Data and analysis; average % core shown in Fig. 5D). Several sequons could not be assigned a glycan (e.g., N135 and N138), as their proximity made it difficult to isolate peptides with only one glycan.

Compared to PV Env trimer glycans, soluble SOSIP trimer glycans are less differentiated, exhibit more variable maturation states and more common sequon skipping (67, 80, 84). A comparison of ‘parent’ trimers and monomeric JR-FL gp120 revealed that glycan types (i.e., high mannose or complex) were similar at many positions (S1 Data and analysis, S8 Fig). However, gp120 monomer glycans were more differentiated at positions N88, N156, N160, N241, perhaps reflecting the greater access to glycan processing enzymes (Fig. 5C). Conversely, glycans N295 and N301 were less mature, which, as for SOSIP, may be because accelerated Env production reduces contact time with glycan processing enzymes (S1 Data and analysis).

### Sequon optimization

Sequon skipping was common in gp120: 8 out of 14 sequons were partially unoccupied (Fig. 5D, S1 Data and analysis). Of these, N156 was most frequently skipped (66.7%), followed by N301 (33.5%) and N362 (8.9%). As we noted above, N156 and N362 were also occasionally skipped in functional trimers, but not to the extent observed for gp120 monomer. Both sequons have a serine at the 3^rd^ position, i.e., NXS (Fig. S1). Since NXT is a more effective substrate for glycan transfer than NXS (84, 105), we made S158T and S364T mutants. Neither mutant affected expression. S158T infectivity was also unchanged, but S364T infectivity was reduced to ∼15%. Glycopeptide LC-MS revealed that S158T mutant glycosylation was broadly similar to the parent, with score changes <+/-5 (Fig. 5C, S1 Data and analysis, S8 Fig). Differences in both directions were observed, e.g., at positions N156 and N301. Maturation states of the complex gp41 glycans also differed, but these are prone to vary, as mentioned above (S1 Data and analysis). Elevated sequon skipping/core glycans at N160 could be a direct consequence of the rare S158T mutation adjacent to this sequon. Significant skipping also occurred at position N339 (Fig. 5D, S1 Data and analysis). Overall, S158T caused some unwanted changes in glycan maturation and skipping.

S364T dramatically increased glycan differentiation at N332 of the gp120 outer domain (Fig. 5C, S8 Fig), as did N301 and N386, to a lesser extent. N262 glycan data was not obtained. Significant sequon skipping occurred at N339 and N637. The N362 glycan is not proximal to N295 and N332, suggesting allosteric effects of this mutant, as for S158T. Both S158 and S364 are well-conserved across strains (S1 Fig), suggesting that they are structurally important. We therefore suggest that sequon optimization be considered only when it does not disturb conserved residue(s).

D197N successfully and completely knocked in the N197 glycan (S1 Data and analysis). (Fig. 5C, S8 Fig). N301 maturation modestly increased. Gp41 glycans varied in complexity (S1 Data and analysis). Unlike the S158T and S364T mutants, N160 and N637 sequons were fully occupied. However, like these other mutants, some skipping occurred at position N339 (Fig. 5D, S1 Data and analysis). Overall, the effects of D197N on other glycan sites were milder than those of S158T and S364T (Fig. 5C, S8 Fig). Modeling suggests that the enhanced N301 glycan maturation is a localized effect (Fig. 5A, S7 Fig, S8 Fig), so it appears that overall trimer conformation is not perturbed.

Sequon-optimized D197N+S199T was inferior to D197N. It only filled the N197 site with glycan to ∼90% and caused dramatic glycan holes elsewhere, most notably at N463 that was ∼91% skipped (S1 Data and analysis). N262 partly toggled to complex. The N301 glycan became immature. These differences help in comparison of the D197N+S199T versus D197N mutant (S1 Data and analysis, S8 Fig). As for the S158T and S364T mutants, distal glycans were affected, suggesting a global ripple effect of the D197N+S199T mutant. Overall, D197N+S199T was not as well-tolerated as D197N, further cautioning against mutations that impact conserved positions (S1 Fig).

T49N successfully added a complex N49 glycan that caused decreased glycan maturation at positions N188, N616, and N637 (Fig. 5C, S1 Data and analysis, S7 Fig, S8 Fig), presumably due to overcrowding and contrasting sharply with the mild effects of D197N. Overall, the reduced gp41 glycan complexity is consistent with S6 Fig. We could not obtain glycopeptide data for N611 that would have given more complete insights into how the N49 glycan improves VRC34 sensitivity. While N611 is not close to the N49 glycan (Fig. 5A), it is possible that smaller glycans at the other gp41 sites provide space for the N611 glycan to move aside for VRC34 binding. Glycan maturation at N301 was also modestly impacted. Our model suggests that some effects are localized (Fig. 5A), but others (e.g., N188) are distal, suggesting a global conformational change, consistent with partial V3 non-nAb sensitivity (Fig. 4, S5C Fig).

N611Q led to increased N262 glycan maturation, decreased N463 glycan maturation and partial skipping at N188 and N339 (Fig. 5D). T49N+N611Q caused reduced N301 maturation (Fig. 5C, S8 Fig) and skipping at positions N339 and N463 (Fig. 5D). Reduced N301 maturation of the single T49N and N611Q mutants appeared to be amplified in the double mutant. However, other effects in the single mutants were absent in the double mutant, suggesting that the N49 knock in may partially compensate for N611 glycan loss. Comparing the double mutant to its component single mutants again highlighted differences at positions N188, N262, N301 and N463 (S8 Fig), although the patterns did not resemble those above, implying unpredictable and subtle effects on trimer folding.

Analysis of N138A+N141A revealed that the N135 glycan is complex (at least in the absence of these neighboring glycans). N135 was not detected on the parent, probably due to its proximity to N138. Significant skipping at N262, N295 was observed, and, to a lesser extent, at N339 (Fig. 5D). The small amount of glycan detected at position N262 was far more mature than on the parent (Fig. 5C). Although this mutant was infectious (Fig. 4A, lane 12), since this glycan is structurally important, its absence could cause some misfolding (100). Glycan maturation differences were also observed at positions N156, N188, N616 and N637.

N138A+N141A Env complexed with CH01 exhibited radical glycan changes at some positions: a shift to high mannose glycans at positions N135 and N188 is consistent with a preference for high mannose glycans to minimize clashes at the binding site (S7 Fig, S8 Fig, Fig. 5A) (13). However, N356 and N463 glycans also increased in high mannose, despite being distal from the CH01 epitope. Intriguingly, glycan N262 became less mature, while glycan N295 was largely complex in CH01-bound sample, despite both being skipped in the unbound sample. Conversely, the N332 glycan exhibited more skipping and glycan N616 was more complex in the CH01-bound sample. These notable findings reveal the presence of glycan species in CH01 complexes that were not detected in the reference sample and *vice versa*. Thus, some glycans exhibit more extensive variability than expected. The differences may reflect the idea that CH01 neutralizes the N138A+N141A mutant to a maximum of only ∼75%, suggesting that it binds only a fraction of trimers where N135 and N188 glycan clashes are minimal, that this fraction of trimers carries other glycan variants that may further improve CH01 binding or that are inextricably linked with the presence of small high mannose glycans at position N135.

Overall, these mutants reveal that outer domain glycans (N156-N339) are prone to maturation changes, while inner domain glycans N88, N356-N448 are largely unchangeable. Sequon skipping was also more common at some outer domain glycan sites, particularly N339. Overall, our findings shed light into the hitherto unknown effects of mutations on local and distal glycans and which changes are well-tolerated or otherwise for potential vaccine use.

#### Final JR-FL mutants

A final set of JR-FL mutants were made to combine and try to improve on the best features so far, starting with the triple V1 glycan deletion mutant in Fig. 4B, lane 13. Overlaying the N611Q mutation improved VRC34 sensitivity as expected, while V2 NAbs were largely unaffected except for a modest loss of PG9 sensitivity (Fig. 4B, lane 32). Previous studies suggested that modifying V3 sequence (106) and an S365V mutant (107) may improve V2 NAb sensitivity. However, trimers mutated with a global V3 consensus sequence (lanl.gov) did not express efficiently, and S365V had little effect (Fig. 4, lanes 33 and 34). This suggests that cognate V2-V3 sequences are important for folding and that the effect of S365V is context-dependent.

Highly basic C-strands may initiate V2 NAb lineages via electrostatic interactions (36, 69, 108). However, a V169R mutant to render the JR-FL C-strand more like many V2-sensitive group 1 strains (Fig. 1) - was misfolded (Fig. 4, lane 17), as described above. A D167N mutant provide another way to increase strand C charge (Fig. 4B, lane 35), as found in some V2-initiating sequences (69). This further increased CH01 sensitivity (Fig. 4B, lane 35, S5A Fig). In fact, it was broadly sensitive to all 4 V2 NAbs and is the most V2-sensitive JR-FL mutant. We mark the lane number of this most effective mutant in red. However, it was also somewhat V3-sensitive (S5C Fig) and PGT145 saturation was somewhat reduced (S5D Fig). Taken together, the increased V3 sensitivity and reduced PGT145 saturation suggests a slightly more ‘open’ trimer apex. Despite its improved CH01-sensitivity, this mutant was still resistant to the CH01 UCA (Fig. S5A). A further increase C-strand charge via D167K mutation reduced V2 sensitivity (S5A Fig and S5D Fig). This is perhaps not surprising, given that D and N are the only permissible residues at this position (S4 Fig).

During V2 NAb ontogeny in natural infection, the C-strand may become more neutral, as the virus attempts to escape NAbs. In turn, V2 NAbs evolve to be less dependent on electrostatic charges and depend more on V2 “anchor” residues (36, 54, 69, 108). To mimic the “escape” phenotype of such “late” viruses, we made an R166K+V169E variant. We also added back the V1 and N611 glycans and modified the FP sequence to the second most common variant (8). Including these changes in boosts could help V2 and FP NAbs evolve to tolerate sequence variations and navigate glycans. However, none of the V2 NAbs neutralized this variant, most likely because V169E eliminates V2 binding completely. However, the array of other mutants in Fig. 4B provide a variety of options to increase V2 stringency in boosts without eliminating V2 sensitivity altogether. Conversely, VRC34 neutralized I515L comparably to other mutants that retain the N611 glycan, suggesting that it tolerates this FP sequence variation.

Finally, we investigated approaches to improve JR-FL Env processing at the lysine rich gp120/gp41 junction (see S1 Text). While we were unable to improve cleavage efficiency by mutation or furin co-transfection, data suggest that a lysine or arginine at position 500 (S1 Fig) should be used as a repair mutation in strains where other residues are present.

### An alternative PV neutralization assay for poorly infectious clones

Our standard neutralization assay uses pNL-LucR-E- and an Env plasmid to make PV for infection of CF2.CD4.CCR5 cells (NL-Luc assay). In this assay, Q23 SOS gp160ΔCT PV infection was low and close to our arbitrary cutoff of 50,000 relative light units (RLUs), where neutralization becomes difficult to distinguish from background. The “gold standard” TZM-bl protocol cannot be used for SOS PV infection, as it involves overlaying cells on virus-antibody mixtures, which is incompatible with our requirement to wash cells after briefly exposing them to 5mM DTT to break the SOS bond of spikes attached to cellular receptors, allowing infection to proceed (103). We therefore sought a different protocol that uses pre-attached cells. We adapted a PV assay previously reported for coronaviruses, in which viral budding is driven by MuLV GagPol and luciferase is carried by plasmid pQC-Fluc (109). PVs made this way mediated elevated infection versus the NL-Luc assay for poorly infectious Q23, WITO and T250 SOS PV (Fig. 6A). However, JR-FL SOS PV infection (already high in the NL-Luc assay), was slightly lower in the pQC-Fluc assay. To check if neutralization sensitivity was impacted by assay differences, we compared PG9 neutralization of the four viruses in both assays. The NL-Luc assay resulted in high error bars compared to the pQC-Fluc assay, most notably for Q23 and WITO (Fig. 6B). Nevertheless, PV PG9 sensitivities were comparable, suggesting that the pQC-Fluc assay is a reasonable substitute whenever infectivity is too low in the NL-Luc assay.

**Figure 6.**
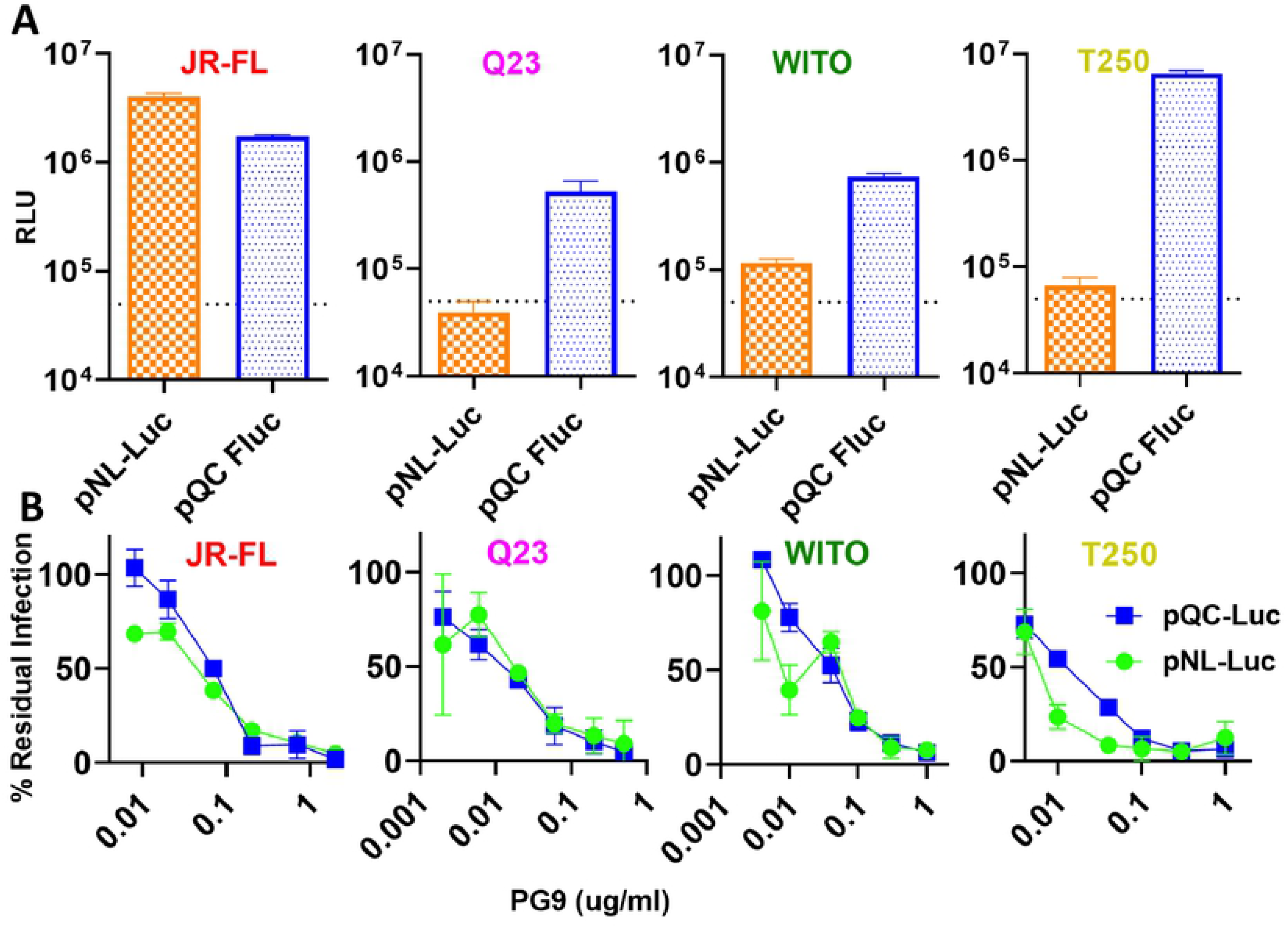
Comparison of NL-Luc and pQC-Fluc assays for HIV pseudovirus infectivity and neutralization sensitivity. A) JR-FL E168K+N189A, Q23 D49N+N611A, WITO and T250 SOS gp160ΔCT PVs, produced using NL-Luc or pQC-Fluc plasmid sets were compared for infection of CF2.CD4 CCR5 cells, assayed as RLUs. The dotted line marks an arbitrary cutoff for infection, below which data become unreliable. B) Comparative PG9 sensitivity of the same PV in both assays to the PG9 mAb.

#### Q23

Q23 is highly CH01 UCA-sensitive (Fig. 2) and expresses well (Fig. 3B-D), so may be ideal for V2 NAb priming. Q23 SOS is also highly V2-sensitive (Fig. 7B, lane 1, S9A Fig). To try to further increase Q23’s V2 sensitivity, we removed the two V1 glycans at positions N133 and N138 alone and together. These mutants reduced infectivity and expression (Fig. 7A, lanes 1-4), but had little effect on V2 NAb or CH01 UCA sensitivities, except that the weak PGT145 sensitivity of the parent virus was lost (Fig. 7B, lanes 1-4, and S9C Fig).

**Figure 7.**
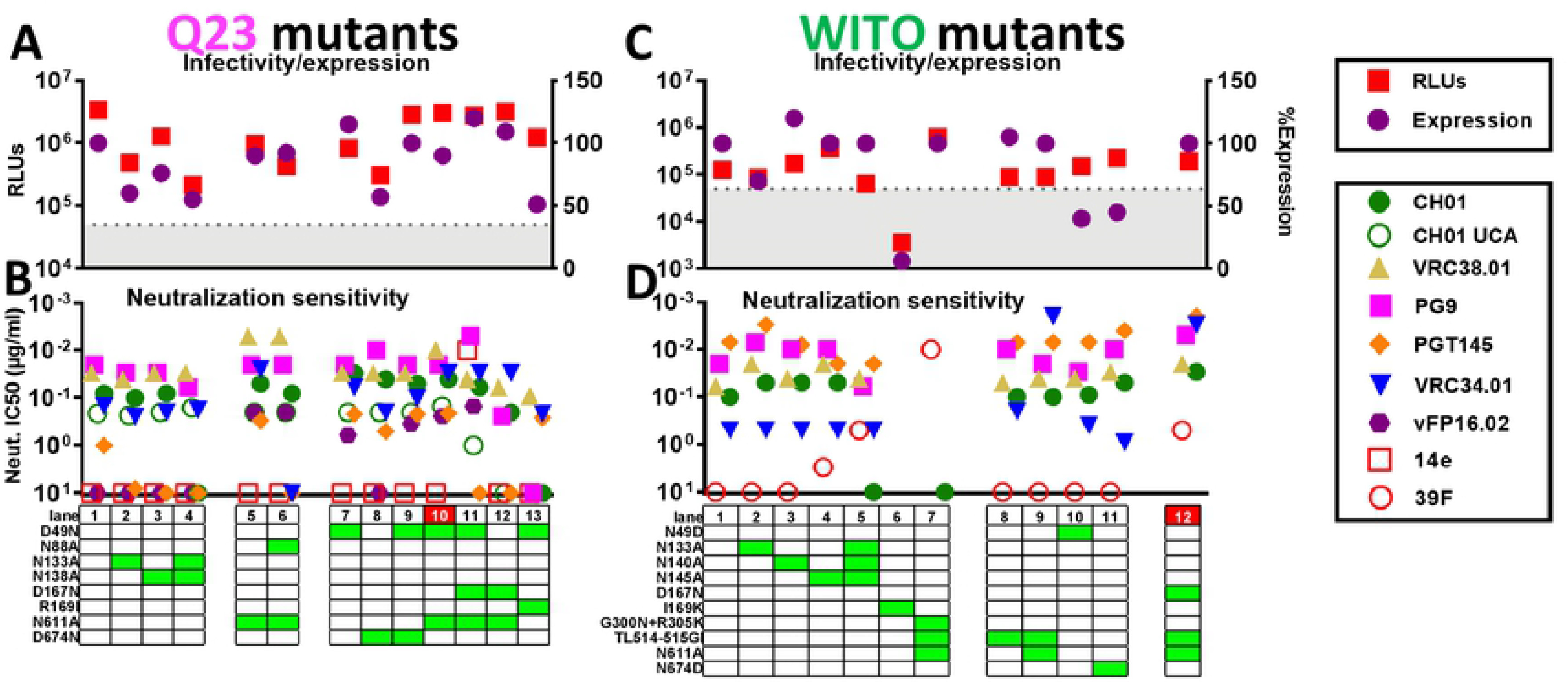
Effects of Q23 and WITO SOS gp160ΔCT mutations on trimer expression, infectivity and NAb sensitivity. Effect of mutations on A) Q23 and C) WITO gp160ΔCT SOS trimer infectivity (by the pQC-Fluc assay) and total Env expression (by SDS-PAGE-Western blot). B) and D) MAb sensitivity.

To try to increase FP NAb sensitivity, we tested the effect of N611A alone and together with N88A. As for JR-FL, N611A improved VRC34 sensitivity. Vaccine-elicited FP NAb vFP16.02 also neutralized this mutant (Fig. 7B, lane 5). When N88A was overlaid, vFP16.02 was still able to neutralize, but VRC34 sensitivity was lost, consistent with VRC34’s requirement for N88 glycan (Fig. 7B, lane 6).

We next examined the effects of knocking in N49 and N674 glycans, as found on well-expressed strains WITO and AC10 (Fig. 1). Considering Q23 trimer’s low total glycan number (Fig. 1, S1 Fig), these additional glycans might assist trimer folding. D49N slightly improved expression, whereas D674N reduced expression (Fig. 7A, lanes 7 and 8). Together, there was no net effect on expression (Fig. 7A, lane 9). Notably, D49N knocked in vFP16.02 sensitivity and increased VRC34 sensitivity (Fig. 7B, compare lanes 1 and 7), as for JR-FL (Fig. 4). PGT145 and CH01 sensitivities were also slightly higher (Fig. 7B, compare lane 1 to lanes 7-9).

D49N+N611A expressed well and was robustly VRC34- and vFP16-sensitive (Fig. 7B, lane 10). CH01 sensitivity was moderately higher than the parent (S9A Fig). 14e saturation also increased, although it did not achieve an IC50, suggesting partial exposure (S9D Fig). CH01 UCA sensitivity was unaffected (Fig. 7B, lane 10, S9B Fig). Overall, this further suggests that the N49 glycan opens the trimer slightly to expose V2 and V3 targets. To try to further increase V2-sensitivity, we overlaid the D167N mutant. However, this showed loss of PGT145, CH01 and CH01 UCA sensitivity, overt V3 sensitivity and poor expression (Fig. 7A and B, lane 11, S9A, B and D Fig). Since N49 and D167N may both modestly increase V3 sensitivity, together they may lead to overt V3 sensitivity. We therefore tested the effects of D167N without D49N. Although V3 resistance was restored, this mutant was still poorly V2-sensitive (Fig. 7B, lane 12).

Considering the negative impact of the V169R mutation on JR-FL (Fig. 4B), we wondered if essentially the reverse mutation, i.e., knocking out Q23’s basic residue by a R169I mutation might improve its expression. However, this was not the case, and sensitivity to V2 and FP NAbs was reduced or eliminated (Fig. 7B compare lanes 7 and 13).

Given Q23’s CH01 UCA sensitivity, we tested if it could also stimulate CH01 UCA expressed on the surface of B cells *ex vivo*. Total (CD19+ B220+) B cells from the spleens of naïve CH01 UCA ‘double knock in’ mice, i.e. expressing both heavy and light chain rearrangements (110) were effectively labeled by WITO SOSIP (S10A Fig). As expected, an anti-IgM Fab2 induced calcium flux. Q23 SOS D49N+N611A VLPs also stimulated *ex vivo* CH01 UCA dKI+ splenic B cells effectively and this result titrated (S10B Fig). GnT1-VLPs induced more robust stimulation, whereas bald VLPs did not stimulate cells (S10B Fig). Thus, Q23 VLPs may be highly effective at priming CH01-like specificities in a vaccine regimen, especially if they are produced in GnT1-cells.

#### WITO

We next attempted to improve WITO, another well-expressed (Fig. 3B), V2-sensitive (Fig. 2) group 1 strain. Above, we saw that CH01 neutralization of WITO gp160ΔCT WT PV was sub-saturating and did not reach an IC50 (Fig. 2). However, using the pQC-Fluc assay to improve WITO SOS PV infection (Fig. 6), we found that like JR-FL SOS PV, WITO SOS PV was neutralized by CH01 with somewhat better saturation, and an IC50 was measurable (Fig. 7D, S11A Fig). We next checked the effects of removing the 3 V1 glycans at positions 133, 140 and 145 alone and together. Expression of N133A was lower, but N140A and N145A expressed like the parent (Fig. 7C, S11B Fig). Sensitivities to multiple V2 NAbs were slightly higher (Fig. 7D, lanes 2-4). However, removing all 3 glycans together reduced CH01 sensitivity and also caused some V3 sensitivity (Fig. 7D, lane 5 and S11A Fig).

An I169K mutation in strand C might improve V2 sensitivity. However, expression and infectivity was poor (Fig. 7C, lane 6, S11B Fig), as with the equivalent JR-FL mutant (Fig. 4A). We attempted to improve apex folding via G300N+R305K mutations at the V3 loop base that may interact with Y173 and Y177 of the V2 loop (104). To accelerate screening, we combined this double mutant with TL514-515GI (FPvar1) and N611A to knock in FP sensitivity. However, this mutant was overtly V3-sensitive and CH01-resistant (Fig. 7D, lane 7, S11A Fig) - essentially the reverse of the desired effect. The same mutant lacking the G300N+R305K (Fig. 7D, lane 9) was not overtly V3-sensitive, suggesting that G300N+R305K causes misfolding. TL514-515GI slightly improved VRC34 sensitivity (Fig. 7D lane 8), and, as expected, combining this with N611A led to a dramatic further increase in VRC34 sensitivity (Fig. 7D, lane 9).

We next tested the effects of *removing* the unusual N49 and N674 glycans that also exist in the well-expressed AC10 strain (Fig. 1, Fig. 3, S1 Fig). Both mutations reduced Env expression (Fig. 7C, lanes 10 and 11, S11B Fig). Analysis of 4,582 M-group Env sequences from the Los Alamos HIV Database reveals that the N49 glycan is ∼10% conserved (S1 Fig) and more common in clade B, moderate in clade D, F1, G, AE and virtually absent elsewhere (S11C Fig). Above, we saw that knocking in the N49 glycan had no effect on JR-FL expression (S6 Fig) and improved Q23 expression slightly (Fig. 7A), suggesting that N49 impacts expression in some, but not all scenarios. The N674 glycan is slightly more prevalent (13% conserved) and is present in 5 of our 17 strains (Fig. 1, S1 Fig). However, none of these other N674 glycan-containing strains were well-expressed, suggesting that the N674 glycan alone does not partition with high expression.

Finally, we combined the FP-immunofocusing mutant TL514-515GI+N611A with D167N to try to create a highly V2- and FP-sensitive combination mutant. This improved WITO sensitivity to multiple V2 NAbs, albeit with a moderate increase in V3 sensitivity (Fig. 7D, lane 12, S11A Fig), similar to the JR-FL D167N mutant (Fig. 4, lane 35, S5A Fig).

#### T250

T250 is a well-expressed group 1 strain. We initially repaired the gp160ΔCT SOS parent, filling glycan holes at positions 276 and 448 and optimizing the gp120/gp41 processing site. These repairs slightly reduced expression and infectivity (Fig. 8A, lanes 1 and 2). Both were PG9-, CH01- and weakly CH01 UCA-sensitive (S12A, B Fig) - the latter being a rare feature so far shared only with Q23. However, the repaired mutant was PGT145-resistant (S12D Fig). Poorly saturating 14e neutralization suggested partially open trimers, so it was unsurprising that the D49N mutant led to overt V3 sensitivity (Fig. 8B, lane 3, S12C Fig). Removing one or both V1 glycans led to a modest increase in CH01 sensitivity, coupled with decreased V3 sensitivity (Fig. 8B, lane 4 and 5, S12A and C). In contrast, D167N lost V2 sensitivity, similar to the D49N mutant, and became overtly V3-sensitive (S12C Fig).

**Figure 8.**
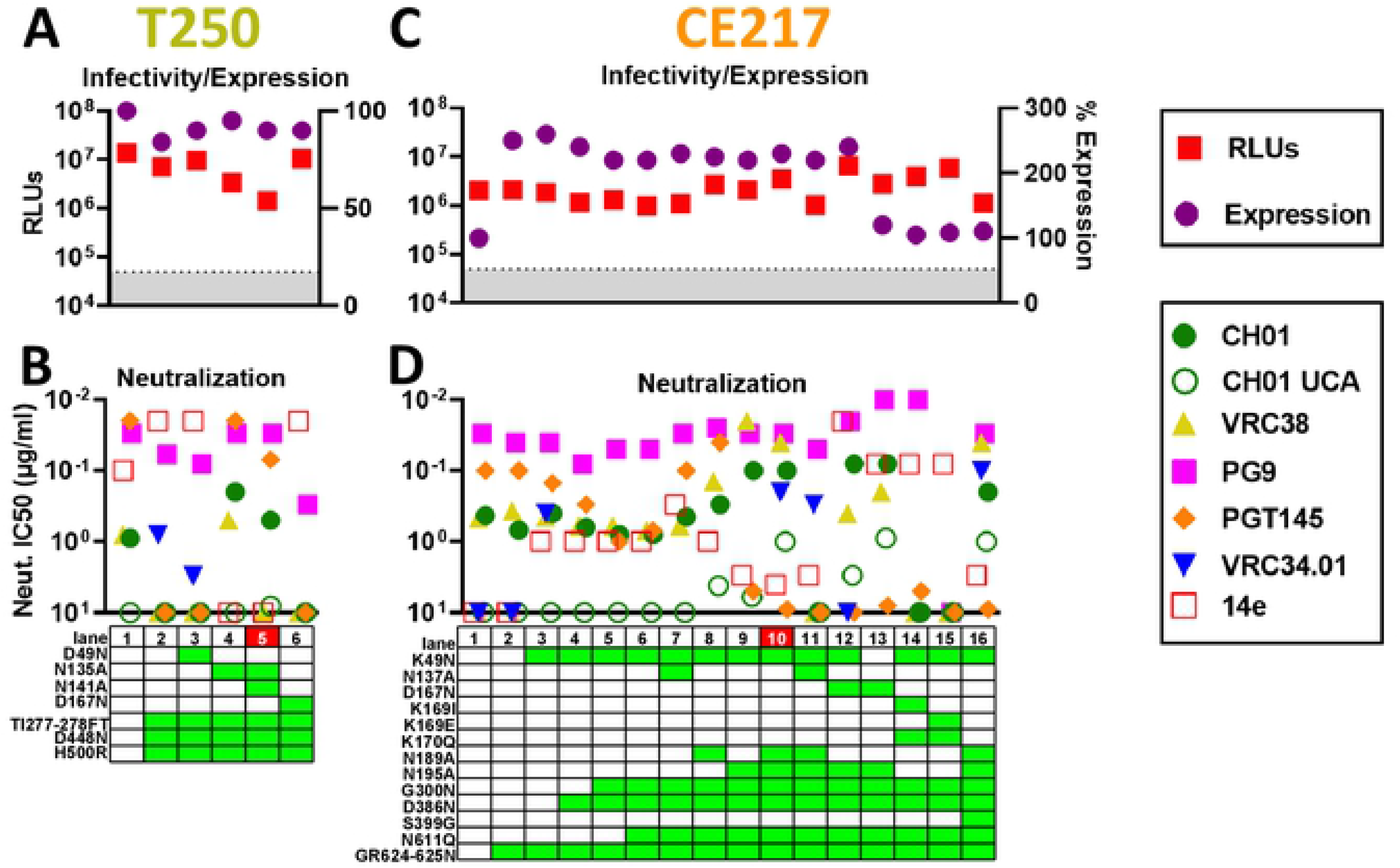
Effects of T250 and CE217 SOS gp160ΔCT mutations on trimer expression, infectivity and NAb sensitivity. Effect of mutations on A) T250 and C) CE217 gp160ΔCT SOS trimer infectivity (by pQC-Fluc and NL-Luc assays, respectively) and expression (by SDS-PAGE-Western blot). B) and D) mAb sensitivity of mutants.

#### CE217

CE217 is among the most V2-sensitive group 1 strains, but is modestly expressed (Fig. 3B-D). Sequence alignments (S1 Fig) reveal an unusual insertion at position 625 (S1 Fig) that could adversely impact gp41 helix folding. We repaired this via a GR624-625N mutation, rendering it more consistent with other strains (S1 Fig). This dramatically improved expression, but did not improve infectivity (Fig. 8C, lanes 1 and 2, S13A Fig). Expression was not further increased by K49N (Fig. 8C, lanes 2 and 3, S13A Fig). D386N to fill in a glycan hole also had little effect, aside from slightly decreased PGT145 and PG9 sensitivity (Fig. 8D, lane 4). G300N (104) further decreased PGT145 neutralization activity (Fig. 8D, lane 5), but had little effect on CH01 and its UCA (S13 Fig B, C). N611Q knocked in VRC34 sensitivity, as expected (Fig. 8D, lane 6). Removing V1 and V2 clashing glycans generally improved V2 sensitivity (Fig. 8D, lanes 7-11). The N195A mutant was highly sensitive to CH01 and VRC38, but lost PGT145 vulnerability. Thus, V2 NAb sensitivities are differentially regulated by N195A (Fig. 8D, compare lanes 6 and 9, S13B, E Fig). N189A and N195A mutants were also measurably susceptible to the CH01 UCA (Fig. 8D, lanes 8 and 9, S13C Fig). Removing both N189A and N195 glycans, however, did not further increase CH01 or VRC38 sensitivity (Fig. 8D, lane 10). Indeed, N137A+N189A+N195A eliminated CH01 and VRC38 sensitivities altogether (Fig. 8D, lane 11). Adding D167N to N195A improved PG9, but not CH01 sensitivity (Fig. 8D, compare lanes 9 and 12). Furthermore, VRC38 and VRC34 sensitivities were reduced and lost, respectively and 14e sensitivity became overt (S13D Fig). Since K49N and D167N mutations may both induce partial V3 sensitivity that become overt when in combination, we reverted N49K. This modestly improved V2 NAb sensitivity and slightly decreased V3 sensitivity that nevertheless remained overt (Fig. 8D, lane 13).

K169I, K169E and K170Q mutants were made to try to reduce V2 sensitivity, for possible late boosting. However, these mutants almost completely eliminated V2 sensitivity, reduced trimer expression and were overtly V3-sensitive (Fig. 8C and D, lanes 14 and 15). Finally, an S399G mutation to nix 397 sequon that overlaps with a more common sequon at position 398 (S1 Fig) did not cause significant changes in mAb sensitivities but reduced expression and infectivity (Fig. 8D, lanes 10 and 16).

#### c1080

c1080 is a clade AE group 1 strain that expresses well (Fig. 3B), and exhibits an average number of glycans and no glycan holes (Fig. 1). Gp160 truncation (gp160ΔCT WT) led to a loss of PG9 sensitivity, which was partly restored by the SOS mutant, albeit with some V3-sensitivity (S14A Fig). D49N markedly improved expression (S14B Fig) but led to overt 14e-sensitivity. This contrasted the modest effect of N49 on the V3 sensitivities of JR-FL and Q23, probably because the SOS parent is already partially V3-sensitive, like T250 SOS parent (S12 Fig). In contrast, N49 had little effect on PG9 or CH01 sensitivities (S14A Fig), again showing that increases in V3 sensitivity induced by mutations can occur without the loss of V2 sensitivities.

The unusual H375 residue of this strain and other AE strains could impact trimer compactness and/or CD4 sensitivity (111). H375S reversion did not improve CH01 sensitivity (S14A Fig). Moreover, like D49N, it was overtly 14e-sensitive (Fig. S14A). These latter mutants are more V3-sensitive than they are to V2 NAbs, suggesting a significant loss of trimer compactness that is incompatible with our goal to immunofocus on V2. Therefore, any further mutants should not be combined with either D49N or H375S. Since the parent virus is partially V3-sensitive, a strategy akin to T250 may be effective (avoiding D167N but removing clashing glycans). However, we did not pursue c1080 further at this point, given that the T250 mutant suffices as a late shot and is also CH01 UCA-sensitive (Fig. 8B, lane 5).

#### AC10

Given our success with JR-FL (Fig. 4), we took a similar strategy with other group 2 strains. The AC10 parent is already PG9- and PGT145-sensitive and lacks a clashing N130 glycan (Fig. 1). The SOS parent and A388T glycan hole-filled mutants both retained PG9 sensitivity and CH01- and resistance to V3 non-nAbs. Data above suggest that “outermost” glycans (i.e. those closer to the N terminus of V1 and C-terminus of V2) may be more prone to V2 clashes (Fig. 4, 7, 8), so we removed these from AC10 first, alone and together with innermost glycans. V1 glycan mutants N137A and N137A+N142A led to modest changes in PGT145 and PG9 sensitivity, but remained CH01- and VRC38-resistant (Fig. 9B, S15A Fig). V2 glycan mutants N185A and N185A+N184A either eliminate an overlapping sequon or both V2 sequons (Fig. 1, S1 Fig), leading to PG9 resistance, but retained PGT145-sensitivity and resistance to 14e, CH01 and VRC38 (Figs. 9B and S15A). Finally, D167N mutant caused increased PG9 sensitivity, detectable CH01 sensitivity, but also partial V3-sensitivity. Although PGT145-sensitivity was intact, it was sub-saturating.

**Figure 9.**
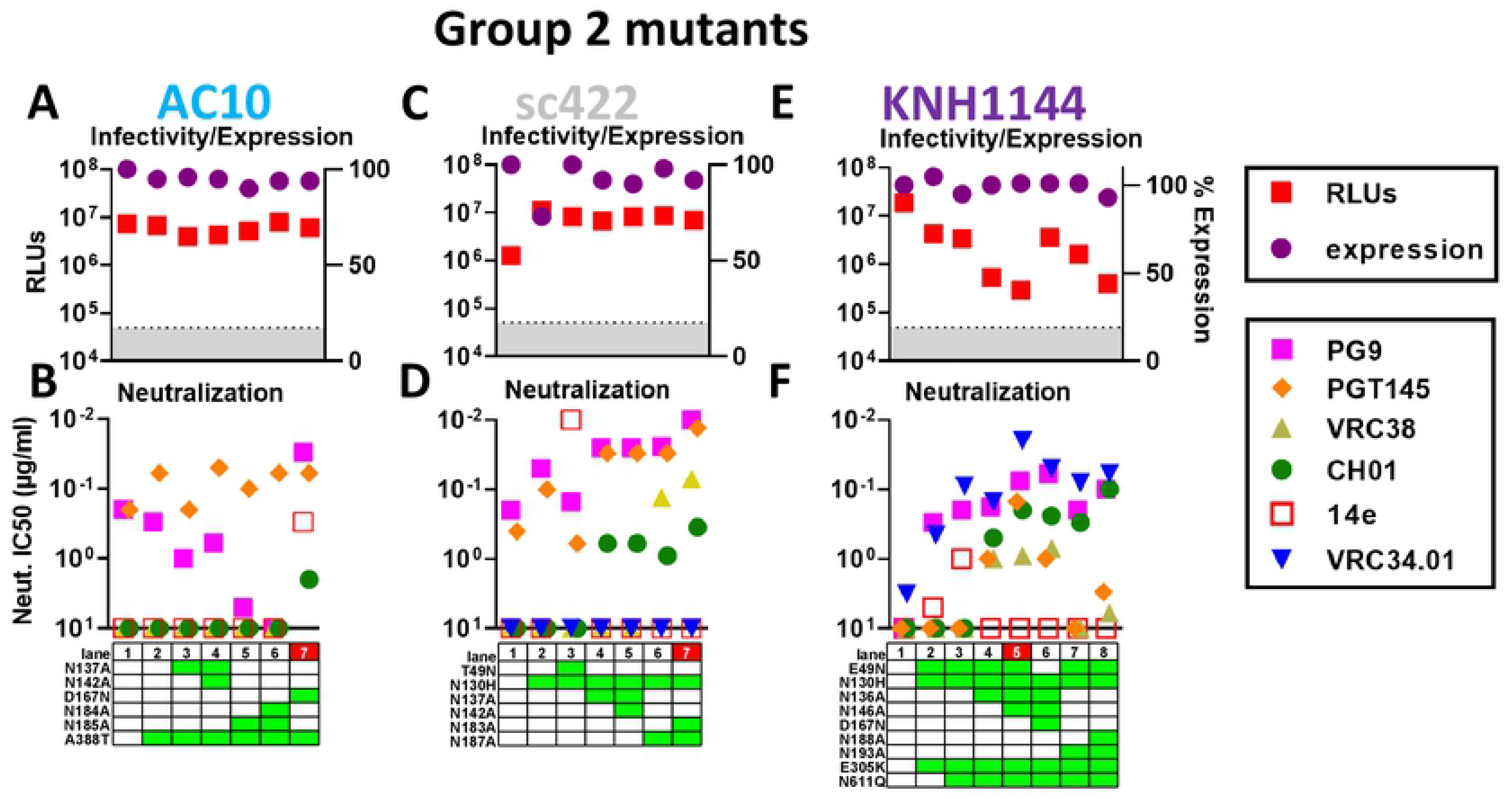
Effects of Group 2 strains AC10, sc422 and KNH1144 SOS gp160ΔCT mutations on trimer expression, infectivity and NAb sensitivity. Effect of mutants on A) AC10, C) sc422 and E) KNH1144 gp160ΔCT SOS trimer infectivity (by pQC-Fluc assay) and expression (by SDS-PAGE-Western blot). B), D) and F) mAb sensitivity of mutants.

#### sc422

sc422 is the best expressed clone in our panel (Fig. 3B-D) and is also PGT145-sensitive (Figs 1 and 2A). Although neither PG9 nor CH01 neutralized the WT parent (Figs. 1 and 2), the SOS mutant was sensitive to both (Fig. 9D), similar to findings with SOS mutants of JR-FL and WITO (S3 Fig). N130H mutation further increased PG9 and PGT145 sensitivity, but CH01 and VRC38 resistance remained intact (Fig. 9D, lane 2, S15B Fig). Knocking in the N49 glycan led to overt V3 sensitivity and reduced PG9 and PGT145 sensitivity (Fig. 9D, lane 3, S15B Fig). Removal of V1 and V2 glycans significantly increased sensitivity to multiple V2 mAbs (Fig. 9D, lanes 4-7 and S15B Fig). Removal of V2 glycans N183 and N187 knocked in VRC38 sensitivity. The N183A+N187A mutant was the most sensitive to multiple V2 NAbs and retained complete V3-resistance.

#### KNH1144

KNH144 is a poorly V2-sensitive group 2 strain. Accordingly, we made several initial mutants in combination: E49N to try to maximize expression, N130H to eliminate a V2 clashing glycan and E305K to try to improve V1V2 packing (104). These changes knocked in PG9- and VRC34-sensitivity and sub-saturating 14e sensitivity (Fig. 9F, lane 2). CH01 sensitivity was also detected (S15C Fig) but did not reach an IC50 (Fig. 9F). PG9 sensitivity may be due to N130H mutation and/or E305K. VRC34 and 14e sensitivities were likely a result of E49N. N611Q improved VRC34 sensitivity, as expected (Fig. 9F, lane 3). Finally, we removed potentially clashing V1 and V2 glycans, starting with those closest to the base of each loop (i.e. N136A and N193A), then double mutants. Both single mutants (N136A and N193A) resulted in detectable CH01 IC50s, albeit sub-saturating (Fig. 9F, lanes 4 and 7). Double mutants (N136A+N146A and N188A+N193A) both improved CH01 IC50s slightly as well as its saturation (Fig. 9F, lanes 5 and 8, S15C Fig). Only the V1 glycan knockouts resulted in moderate VRC38 and PGT145 sensitivity (Fig. 9F, lanes 4 and 5). We attempted to increase the sensitivity of the mutant in Fig. 9F, lane 5 by a D167N mutation and reverting the N49 glycan. This had surprisingly modest impact on V2 sensitivity, except for a loss of PGT145 sensitivity. VRC34 neutralization was also reduced, suggesting that the E49N and N611Q mutations both assist VRC34 sensitivity in this case (Fig. 9F, lane 5 and 6), unlike that in JR-FL (Fig. 4).

#### 6101

Of the group 2 strains, 6101 is the poorest expressing (Fig. 3B-D) and also lacks V2- sensitivity (Fig. 1). Nevertheless, we attempted 6101 repair using combinations of mutations including T49N, N130H, D160N, DK166-167RD, T171K, D177Y, GG269-269E, 355G to optimize V2 sensitivity and resolve insertions and deletions. Although some mutants were quite infectious with RLUs >500,000, neutralization data was difficult to interpret so we did not pursue this strain further.

### Other engineering approaches

In addition to the engineering approaches above, we evaluated many others, exemplified in S2 Text.

### A quarter of particles produced from transfected cells carry surface Env trimers

We previously found that codon-optimized MuLV Gag drives higher yields of Env trimer on VLPs, compared to pNL-LucR-E- (23). Electron microscopy showed that some particles bear surface spikes (77, 112), although “bald” particles with no spikes are also common. We were concerned that MuLV Gag-induced budding might outpace surface Env expression, decreasing the proportion of “Env+” VLPs and/or spike density.

To investigate the Gag-dependency of particle trimer expression, we co-transfected a fixed amount of WITO SOS gp160ΔCT with 10-fold decreasing Gag doses. We also transfected WITO SOS gp160ΔCT alone to assay spontaneous (i.e. endogenous) Env+ particles. Another sample was generated by transfecting a high dose of Gag alone. Supernatants were filtered and 1000x concentrated. As expected, higher doses of MuLV Gag drove production of more particulate Env (Fig. 10A). However, Env was detected even when 1,000-fold less Gag or no Gag was co-transfected (Fig. 10A, compare lanes 1, 4 and 5). Env was not detected in the Gag only sample (Fig. 10A, lane 6).

**Figure 10.**
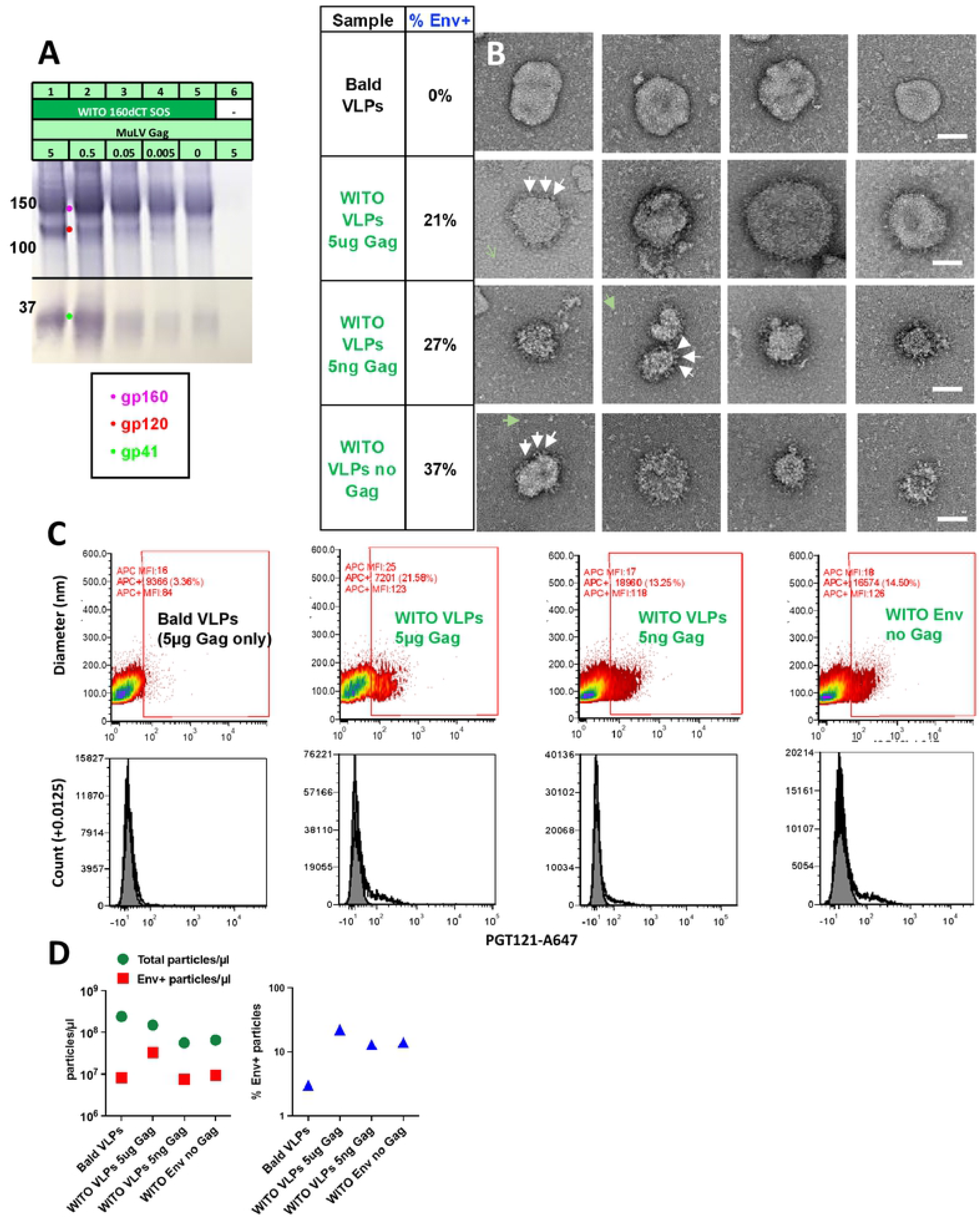
A quarter of particles from transfections with Env plasmid carry surface Env. 293T cells were transfected with WITO SOS gp160ΔCT and/or MuLV Gag, as indicated in part A). Supernatants were precleared, filtered and 1,000-fold concentrated. Samples were probed by A) SDS-PAGE-Western blot, B) negative stain EM (scale bars are 50nM, white arrows point to candidate Env trimers and green arrows point to candidate dissociated spikes, and C) Single vesicle flow cytometry. Upper panels show particle diameters and fluorescence intensities of samples stained with Alexa-647-labeled PGT121. In the lower panel, we show total particle counts versus Alexa-647 fluorescence. D) Total particle counts and Env+ particle counts per µl of the samples indicated (left) and % Env+ particles as a proportion of total particles (right). Raw vFC data files and data analysis layouts have been deposited in Flowrepository (flowrepository.org; see S1 Table)

In negative stain EM images (Fig. 10B), Gag only “bald” VLPs (Fig. 10B, top row) provided a reference to help identify Env spikes on other samples. WITO Env-transfected samples (Fig. 10B, rows 2-4) revealed particles with surface structures not detected on bald VLPs that we infer as Env spikes. These putative spikes do not adopt clear propellor-like structures, perhaps because, unlike rigidified near native soluble trimers, native, membrane trimers are flexible and sample different conformations that are harder to resolve.

In addition to spike-bearing particles (Fig. 10B, rows 2-4), bald particles like those in the top row were also prevalent in all the other samples. Counting Env+ and Env-particles in each sample revealed that approximately a quarter to a third were Env+. There was a trend towards a greater proportion of Env+ particles in samples made with little or no Gag. Thus, our concern that efficient Gag-induced particle production might lead to an overwhelming proportion of bald VLPs appears to be unfounded. We noted the presence of some debris, indicated by green arrows in Fig. 10B, rows 2-4, that may be detached spikes, that may dissociate due to VLPs collapsing during the process of negative staining. Finally, the size of particles varied. Many particles were approximately 100nm diameter, although some were much larger, up to ∼300nm in diameter. It is not clear if the latter are Gag-driven particles or Gag-independent extracellular vesicles.

We next analyzed the same VLP samples by single vesicle flow cytometry (vFC™, Cellarcus Biosciences), which uses a fluorogenic membrane probe to detect and size vesicles, and fluorescent mAbs to measure vesicle surface cargo by immunofluorescence (113, 114). Using this method with Alexa647-labeled PGT121 revealed a modest, but consistent proportion of Env+ particles bound PGT121 (13.25-21.58%) in samples transfected with WITO Env (Fig. 10C, columns 2-4, Fig. 10D, right plot). In contrast, VLPs from cells expressing Gag but no Env (“bald VLPs”) showed only background PGT121 binding (Fig. 10C, first column). Transfecting with Gag alone yielded the highest particle count of 2.4×10^8^/µl of sample (Fig. 10D). Co-transfecting Env reduced the particle count by ∼60% to 1.5 x 10^8^/µl (Fig. 10D, left). Transfecting with only 5ng or no Gag reduced particle counts a further 2-fold (Fig. 10D, left). We can infer that ∼25% of particles in these samples form spontaneously and are Gag-independent, putting a new perspective on the increased Env output when transfecting with high amounts of Gag versus no Gag (Fig. 10A, compare lanes 1 and 5). Essentially, transfecting with a high dose Gag increases particle numbers by only ∼4-fold with a concomitant increase in Env, as particle production is already high even with no Gag transfection. Overall, this data agrees quite well with the EM data in that particles are 13-37% Env+ and the proportion of Env+ particles does not change much with Gag co-transfection, which is perhaps not surprising, if Gag only raises above spontaneous particle production levels by about 4-fold.

The single vesicle flow cytometry analysis revealed other pertinent information. The x axis in Fig. 10C (PGT121 binding fluorescence) is calibrated in units of mAbs/vesicle to allow estimation of spike numbers per particle. Bald particles (Gag-only) have a median autofluorescence equivalent to 16 Ab molecules (Fig. 10C, left column), with Env transfection producing PGT121+ vesicles (above a threshold of ∼50 molecules) with a mean of ∼125 Ab molecules/vesicle. PGT121 may bind up to 3 molecules per Env trimer. Thus, extrapolating from the number of mAbs bound per vesicle to number of spikes per vesicle involves some uncertainty. Subtracting autofluorescence from the ∼125 PGT121 mAbs/particle lowers the estimate to ∼109 PGT121 molecules per particle. If 3 PGT121 molecules can bind per trimer at saturation, this suggests a spike density of ∼30-40 trimers per particle. This aligns quite well with our earlier estimate of 27 spikes per particle for our early JR-FL VLPs (77). There was not a clear difference in the extent of particle PGT121 binding between samples (Fig. 10C), suggesting that spike density does not vary, consistent with the finding above that the proportion of Env+ particles also does not change drastically with Gag co-transfection. Finally, single vesicle flow cytometry also revealed that most particles were between 70-180nm in diameter, centered around 120nm, but some particles appeared to be much larger, up to ∼300nm, in agreement with EM data (Fig. 10B).

## Discussion

In an effort to increase the breadth of VLP vaccine-induced NAbs, we sought a diverse panel of V2- and FP-sensitive membrane trimers. For new strains to be adopted for vaccine use, we insisted they each satisfy 4 key criteria. First, that they are well-expressed, like our JR-FL prototype. Second that they are high quality, i.e., with sequons fully occupied, gp120/gp41 well-processed and not overtly V3-sensitive (partial V3 sensitivity is acceptable, as long as it is not greater than V2 sensitivity). Third, that, collectively, they cover a range of sensitivities to desired target(s), ranging from acute, UCA-triggering to that of typical transmitted isolates. Fourth, that they are functional, allowing us to monitor immunogens and vaccine sera using PVs made with the *same* trimers.

Our exhaustive engineering efforts are summarized in S16 Fig. At completion, we identified 7 trimers from 17 initial strains that each satisfy our criteria (S16 Fig, bottom row). Modified Q23, T250 and CE217 trimers could be used in priming, boosting with modified WITO, JR-FL, sc422 and KNH1144 trimers. Given our stiff selection criteria, the remaining clones did not ‘make the cut’ for a variety of reasons: poor expression (CAP45, KER2018, CNE58, BB201, 001428, X2278), poor infectivity (CM244, 6101), V3 sensitivity (c1080) and insufficient V2-sensitivity (AC10).

Ultimately, our efforts did not reveal a “magic formula” for identifying useful trimers. The complexities that govern trimer phenotypes in our criteria defy prediction by Env sequence alignments. Nevertheless, to accelerate the analysis of clones in the latter part of this project, we began by initially filling in glycan holes and “repairing” insertions, deletions and overlapping sequons - thus correcting “errors of nature”. Overall, there appears to be no universal strategy to identify desirable clones. That said, we can rule out numerous unhelpful approaches. Below, we consider the merits of each modification (S16 Fig).

### Gp160ΔCT/SOS

As for our JR-FL prototype, ΔCT improved expression of other trimers. Although ΔCT removes retention signals (115), our observation that C-terminal tags reduce trimer expression (S2 Text), suggests that the weaker expression of Envs with cytoplasmic tails is sequence-independent. Our default ΔCT mutant leaves a 3AA cytoplasmic tail (S1 Fig); further truncation did not improve trimer expression. Also like our JR-FL prototype, SOS mutants of various strains improved trimer expression, ostensibly by preventing gp120 shedding.

Most gp160ΔCT SOS trimers were functional in PV assays. Native trimers are flexible: apex folding is influenced by the V1V2 loops that sit atop and mask the V3 loop on the resting spike (95). Our data reveal that many mutations can impact recognition by V2 NAbs and V3 Abs, including many mutations distal from the apex. Although gp41 truncation can lead to overt V3 sensitivity (115), its effect on our trimers was modest, as was SOS (102). Thus, SOS gp160dCT trimers exhibited relevant, V3-resistant phenotypes.

Our findings are in line with the idea that the trimer apex exists in a variety of states, ranging from fully closed and V2-sensitive to fully open, overtly V3-sensitive, and V2-resistant. Trimers from different strains exhibit substantial variation in fine apex topographies (12). Indeed, some SOS gp160ΔCT trimers were partially V3-sensitive (e.g., T250), while others were not (e.g., Q23). Some trimers remained largely V3-resistant in the face of N49 and D167N mutations. In contrast, partly V3-sensitive T250 was prone to become overtly V3-sensitive and therefore had to be dealt with more carefully (S12 Fig). Perhaps the most striking example of how trimer conformations change with mutation was for CE217, where PGT145-sensitivity was lost in the most CH01-sensitive mutants that also gained some V3-sensitivity, then CH01 sensitivity was then lost as V3 sensitivity became overt with more mutations (Fig. 8D). Thus trimers sample a number of conformations, including intermediates that are partially V3-sensitive but retain some V1V2 NAb sensitivity (107). In some cases, the loosening of the V2 apex is beneficial. For example, SOS mutation improved CH01 saturation of JR-FL and WITO (S3 Fig and S11 Fig). However, in other cases, gp160ΔCT SOS mutants were *less* CH01-sensitive than the gp160 WT PV (Q23, T250 and CE217). Thus, the effect of SOS on NAb sensitivity depends on the apex conformation of parent trimers. Notably, PGT145 sensitivity of gp160ΔCT SOS trimers was somewhat reduced in all cases, as neutralization depends on a tightly folded apex that is slightly perturbed by both the SOS and ΔCT mutations. Overall, our findings are consistent with a slightly activated SOS trimer state (116) that is V2-sensitive and not overtly V3-sensitive.

As in our previous studies, ∼10% of Env mutants were overtly sensitive to V3 mAbs (e.g., A328G in (35)), with IC90s <0.01µg/ml, suggesting “splayed” trimers in which the V1V2-V3 interactions of the apex are lost. In many cases, these mutants exhibit reduced expression and infectivity. Given the frequency of these misfolded mutants, it is not surprising that they are well-documented in other studies (3, 34, 104, 117–120). While it may be reasonably inferred that mutants causing this phenotype suggest a role of the mutated amino acid(s) in maintaining quaternary trimer interactions, this is not necessarily the case. Instead, it may simply be that the mutant disturbs trimer quaternary interactions. This latter appears to be the case for a quartet of V3 and V2 mutations (104) that we hoped might tighten the apex, but that instead caused overt V3 sensitivity in WITO (S11 Fig), while having little apparent effect on CE217 and KNH1144. Thus, while mutations throughout the trimer can adversely impact proper folding, attempts to “close” partially open trimers face a complex and nuanced challenge to identify the residues that rendered the trimer open in the first place. For example, in the case of T250, the V1 glycans clearly limit V2 apex folding/V2 NAb sensitivity, but had less effects in other contexts, e.g. WITO, where other factors limit V2 sensitivity.

### Repair mutations

Although ΔCT and SOS mutations consistently improved expression, this was insufficient to move several group 1 strains forward. Improving expression may depend on finding the factor(s) that regulate trimer expression, which may well differ between strains. We considered a variety of possible “repairs”, including eliminating insertions, deletions and overlapping sequons, filling glycan holes, gp160 cleavage site optimization, and knocking in N49 and N674 glycans. We also made a large number of point mutants, including many reported previously (33, 88, 95, 106, 107). Overwhelmingly, these repairs did not markedly improve trimer expression, infection or NAb sensitivity, suggesting that none of them address the underlying reasons for poor membrane trimer expression. Thus, while these repair strategies may be effective in some formats, they are not broadly effective like ΔCT and SOS mutations (96).

Nevertheless, in a few cases, repairs were helpful. For example, CE217 expression improved upon deletion of an insertion in its gp41 coiled coil. Another example was that removing the overlapping N189 sequon in JR-FL improves V2-sensitivity (Fig. 4). In some cases, adding an N49 glycan improved trimer expression (WITO, c1080) but not in others (JR-FL etc). How might the N49 glycan impact trimer expression? Since glycans are added concomitantly with Env translation and slow Env folding kinetics (121), the early addition of N49 glycan may add structure to promote folding.

Other ineffective attempts to improve expression included modified signal peptides and codon optimization. Different expression plasmids or lentiviral vectors were also unable to improve expression consistently. This suggests that transcription is not a major bottleneck for membrane Env expression. Moreover, making lentiviral cell lines is quite labor-intensive and eliminates the flexibility afforded by switching Env plasmids when making VLPs by transfection.

Domain swaps offer the potential benefit of a well-expressed Env as a scaffold for immunogenic domains. Previously, various TM swaps improved Env expression >10 fold (93). However, in our hands, they had little effect on membrane trimer expression (see example in S2 Text). The discrepancy may be due to the fact that, in the previous study, uncleaved Envs were expressed in insect cells that impart only paucimannose glycans. This implies that the benefit of TM swaps is context-dependent.

In our hands, V1V2 and gp120 chimeras were overtly V3-sensitive or non-functional. However, these strategies were more successful for SOSIP (22, 39, 90). The difference may hinge on the use of rigidifying disulfides like DS-SOSIP that prevent misfolding. Conversely, functional trimers are flexible, increasing the potential for engraftments to perturb trimer folding, for example, due to incompatibilities between a new V1V2 loop with the V3 loop of the scaffold.

Our efforts to improve gp120/gp41 processing were also not successful. Non-basic residues at position 500 may reduce processing (S1 Text), but reverting H500R (in combination with other mutations) did not improve T250 infectivity or expression. Although SOSIP processing benefit from the “R6” mutation (122) and furin co-expression (123), both of these approaches led to reduced membrane trimer expression and infectivity. This may be because the furin site is largely inaccessible on membrane trimers, regardless of enzyme or substrate sensitivity. Indeed, JR-FL membrane trimers were among the most effectively processed, while those of many other strains remain predominantly uncleaved.

### SOSIP vs VLP Envs

The scarcity of well-expressed membrane trimer strains is perhaps analogous to the problem of identifying soluble trimers, except that the nature of the challenge differs. Our efforts revealed that mutants with valuable effects soluble trimers were, aside from the SOS mutant, unhelpful in the context of membrane trimers. This underlines the extensive differences between these two forms of Env that present different challenges. Indeed, different strains make better prototypes in the two formats, e.g., JR-FL for membrane trimers and BG505 for SOSIP. Thus, while expression is a problem for membrane Envs, it is not a problem for soluble trimers. Conversely, while a key challenge for SOSIP is rendering them to be closed trimers, this is generally not a problem for membrane trimers, as their membrane context prevents them from naturally adopting a triggered conformation but requires I559P and other mutations in soluble format.

Another challenge was that we insisted that selected membrane trimers be functional, so that immunogens and vaccine sera can be appraised using the same trimers in relevant PV assays, thus empowering our approach. This is a potential advantage over soluble trimers, where appraisal is limited to binding assays that may not track perfectly with neutralization. We were excited to observe that SOS PVs were functional for several other strains aside from our JR-FL prototype. However, we were not surprised that infectious counts were lower, as for JR-FL SOS PVs. This presented a trade-off between deciding either select WT trimers because they are more functional or selecting SOS trimers because they express better. However, the pQC-Fluc assay dramatically improved infection sensitivity so that all strains were useable. While we do not know why the pQC-Fluc assay improves infectious counts, we suspect it is because the MuLV GagPol plasmid drives higher levels of particle budding than the subgenomic pNL-Luc plasmid (23). Whatever the reason, this assay was decisive in allowing us to carry forward membrane SOS trimers that we would have otherwise discarded. The pQC-Fluc assay may thus turn out to be useful more generally for making PV of clones that exhibit poor infectious counts.

### Summary of expression attempts

Overall, membrane trimer expression appears to depend on many factors, rendering it difficult to identify and resolve expression bottlenecks. Thus, it is important to begin with clones that at least express modest levels of Env. One strategy we did not attempt but that may be helpful would be to screen for highly expressed clonal relatives of V2-sensitive (group 1) strains in Genbank (108). However even if successful, such clones may not satisfy our other criteria and may still require rounds of mutation and screening.

### Sequon skipping and optimization

Glycopeptide analysis revealed that sequon skipping is usually limited for membrane trimers, contrasting with monomeric gp120 and soluble trimers (67, 79, 80, 84, 124). The resulting epigenetic glycan holes are problematic because they induce non-NAbs that could distract from immunofocusing strategies (9, 84).

Incomplete N160 sequon occupation could be problematic for our goals. However, we only observed N160 skipping in one parent sample and the S158T mutant. The latter is consistent with a previous study in which S158T caused N160 skipping in BG505 SOSIP (84). Thus, it follows that when two sequons are closely juxtaposed (a 1 amino acid gap in this case), reducing the efficiency of the first site can increase occupancy of the second. This may explain why S158 heavily predominates over T158 in natural isolates. Similarly, sequon optimization mutants S364T and D197N+S199T were unhelpful. Moreover, in both cases, skipping increased at various distal sites. Overall, sequon optimization has little benefit for membrane Env where skipping is uncommon and may adversely affect folding.

Glycopeptide analysis provided insights into the global effects of toggling glycans on the maturation of other glycans. Revealingly, knock in of D197N was far better tolerated than that of T49N. This was manifested in the T49N mutant’s dramatic changes in glycan scores at various positions, as well as sequon skipping at N195 and in the core glycan at N625. While it might be expected that knock in of the N49 glycan would decrease the maturation of neighboring glycans, e.g. N276 and N637, it affected distal glycans. Furthermore, paradoxically, some glycans became *more* differentiated. This suggests that N49 knock in influences the network of closely spaced glycans on the trimer surface with variable and sometimes far reaching effects that are not easily understood by rigid models of trimer structure. At the same time, N49 modestly affects trimer conformation, as evidenced by V3 exposure. These findings again resemble those for BG505 SOSIP, where filling the relatively conserved N241 glycan hole, like N197, was well-tolerated, while filling the less conserved N289 site, like N49 incurred marked global effects on the glycan network (12, 80, 125–127). The effects of glycan knock in can differ between SOSIP and membrane trimers. Thus, for SOSIP, the presence of N197 glycan decreased N156 and N160 glycan trimming, but not for JR-FL membrane trimers, despite the proximity of N197 to N156 and N160 at the trimer apex (Fig 5A).

Removing N611, a well-conserved glycan caused similar ripples in glycan maturation and sequon skipping like N49 knock in, consistent with a role in maintaining trimer architecture. However, the particular glycans affected differed. Notably, some distal glycans became *less* mature, despite the reduced overall glycan count, again suggesting perturbation of the trimer glycan network. The N138A+N141A exhibited a similar phenotype, even though these are protecting glycans and not structural glycans.

Considering sum of the effects of mutants in Fig. 5C, sequon skipping in JR-FL membrane trimers is concentrated in the outer domain of gp120, between positions 156 and 339 and rarely occurs elsewhere. In contrast, sequon skipping is quite common throughout SOSIP trimers, including position N611 of gp41. The outer domain of membrane trimers is also more prone to glycan score changes. These findings conjure a scenario where exposed glycans are more subject to change, whereas buried structural glycans are more consistent.

We speculated that high glycan numbers and/or lack of glycan holes (gaps in structural glycans) might drive folding and associate with high expression. However, glycan toggling in either direction did not consistently impact expression. That said, as covered above, adding the N49 glycan did improve expression for some strains. Furthermore, glycan removal in several cases reduced expression. For example, removing V1 glycans reduced expression of JR-FL, Q23 and WITO trimers. It may be that glycan content is a delicate balance of sufficient glycans to drive folding, but not so many that glycan overcrowding occurs, which could result in high mannose glycan bias (13), sequon skipping or a reduction in folding kinetics.

Overall, the varied effects of glycan toggling on proximal and distal glycans provides reason for caution. For example, a glycan hole that increases V2 apex sensitivity will only be effective if it does not open up other unwanted glycan holes. On the other hand, if “off-target” holes differ between successive vaccine shots, this could limit the problem.

Our mutant screening efforts paradoxically suggest that V1V2 glycans further from the V2 apex in the linear sequence often mediate clashes with V2 NAbs. Thus, for JR-FL, removing N189 and N135 glycans allow saturating CH01 neutralization. The greater effect of N135 glycan knockout compared to N138 or N141 is, however, consistent with its closer structural proximity to the apex (Fig. 5A). Eliminating the outermost glycans also had more benefit for CE217. The same may be true for T250 and KNH1144, although in these cases, to accelerate the discovery efforts, we combined V1 and V2 glycan knock outs, precluding a clear comparison of single glycan knockouts. Furthermore, the N130 glycan is absent in V2-sensitive strains and its removal from group 2 strains KNH1144 and sc422 improved their V2 sensitivities. However, as mentioned above, some V1V2 glycans have an important structural role and can impact trimer expression. For JR-FL, knocking the N197 glycan did not appreciably impact either expression or V2 sensitivity, although toggling this glycan can impact V2 sensitivity of other strains (128).

Given the preference of CH01 for GnT1-virus (Fig. 2, (13)), it was no surprise that CH01 selected for small high mannose glycan at N135 to minimize clashes. This kind of selective binding to glycovariants has been reported previously (84). What we did not expect is that glycans at other sites to also be far less mature. Although this could suggest that CH01 recovered an earlier “high mannose” trimer glycoform, not all sequons were affected. For example, the N616 glycan was *more* mature. It could be that glycan maturation at different sites is co-dependent, via glycan network effects. If so, however, how can we reconcile the rarity and indeed total absence of glycoforms at several sites in the corresponding uncomplexed parent? Considering that CH01 substantially neutralizes the N138A+N141A mutant, we would expect the trimer glycoform it binds to register prominently amid the trimer glycovariants that constitute the parent sample. However, this assumes that all glycoforms are equally infectious, which may not in fact be the case. Indeed, a significant fraction of JR-FL trimers remains uncleaved and therefore non-functional. However, our glycopeptide analysis provides data on the *total* Env on VLPs, regardless of gp120/gp41 processing. Further analysis of mAb-complexed trimers may help us to dissect this issue.

### Modifying the C strand

V2 bNAbs typically bind one-per-spike (62, 95). C strand charge is at a premium during the early ontogeny of V2 NAbs. Therefore, we engineered trimers to increase C-strand’s charge (Fig. 1), taking care to use substitutions that are acceptable based on sequence alignments (S4 Fig). Conserved K/R residues at positions 166, 168, 169 and 171 are important for V2 NAb binding. Of these, 166 and 171 were present in all but 6101. Our attempts to fix the C-strand and other repairs in this strain, however, did not result in sufficiently functional trimers. K/R168 was present in all but JR-FL, where E168K was an effective knock in. D167N, as found in V2 NAb initiating clones consistently improved V2 sensitivity, albeit with a concomitant increase in V3 sensitivity.

Our attempts to mutate residue 169 to a basic residue in the JR-FL and WITO strains both resulted in low infectivity and expression. This was surprising, considering that V169R mutation of ADA improves its V2 sensitivity and decreased V3 sensitivity (95). Thus, the benefits of knocking in a basic residue at position 169 is context-specific. The reverse mutation K169I/E reduced expression and increased V3 sensitivity in CE217. This suggests a role of residue 169 in folding and does not rule out the possibility that more conservative K169R substitutions may improve V2 binding (33).

### Biophysical analysis of VLPs

We investigated the basis for the dominant proportion of bald particles over Env+ particles in the hope that new information might illuminate ways to improve VLP quality and possibly spike density, which may assist NAb priming. In the past, flow cytometric measurement of individual particles has been challenging. However, new state-of-the-art methods supported by a set of accepted guidelines are now enabling quantitative and reproducible measurement of individual extracellular vesicles and their molecular cargo (129). Our findings raise several questions:

***1) Why is particle production so high, even in the absence of Gag transfection?*** The constitutive release of extracellular vesicles (EVs) 293T cells (113, 114) provide a background of particles, among which Gag-or Env-bearing particles are also released. Some particle production may be driven by Env alone, given that Env’s heavy glycosylation promotes ER stress. This may explain why VLPs invariably contain uncleaved gp160, as Env released under stress bypasses furin processing. Some particles may arise from spontaneous budding of endogenous Gag in the 293T cell genome, e.g. HERV. Finally, some particles may derive from FBS that carries ubiquitous vesicles. After transfection, cells are washed in PBS and replaced with 1% FBS medium a day later. Thus, FBS-derived vesicles are likely to co-purify with VLPs.

***2) Why are only ∼25% particles Env+?*** Bald particles may be almost entirely FBS-derived. This explanation is weakly supported by the observation that co-transfecting high doses of Gag+Env leads to higher total particle production than Env alone and a higher total number and fraction of Env+ particles (Fig. 10D, right lanes 2 and 4). This suggests that Env+ particles are produced amid a constant background of FBS vesicles. However, since Gag transfection only increases particle production by 2-fold compared to Env only transfection, hard interpretations are difficult.

***3) Is there a way to purify Env+ particles?*** Immunocapture might work but eluting them may be problematic. If Env-EVs are higher in some other marker (i.e. CD81) because they bud differently or originate from FBS, they could be fractionated.

### How to put SHPB regimens together?

Now that we have identified 7 diverse, well-expressed, V2-sensitive and functional membrane trimers, how will we use them in vaccine studies? A first step will be to fix the FP sequence of all clones to fusion peptide variant 1 (FP8var1) so that we can begin investigating the immunofocusing on this site at the same time as the V2. The most CH01 UCA-sensitive Q23 mutant would be the ideal prime. T250 and CE217 that are also slightly CH01 UCA-sensitive group 1 strains would also be useful in early shots. WITO and the 3 group 2 strains could be used as boosts. It may be useful to prime with D167N+N611Q mutant trimers and/or where clashing glycans are removed, and then gradually reverse these modifications in boosts, perhaps using N49 knock in as an intermediate FP-sensitive boost, after reinstating N611 glycan.

Also in boosts, strand C could be varied to render it more neutral, mimicking waves of diversity in V2 NAb ontogeny in natural infection (24, 36, 69, 108). Thus, by decreasing CH01-sensitivity in boosts, NAbs may gradually develop an ability to navigate glycans and sequence diversity, while gaining N88 contacts for FP NAbs and N160 contacts for V2 NAbs with increasing dependency on conserved anchor residues within strand C. The V3 occlusion and high sequon occupation of most trimers should help avoid inducing non-NAbs to the V3 loop and glycan holes, as long as these imperfections are limited to single shots, the potential for distraction should be limited.

The above sketches of possible SHPB regimens resemble those attempted previously (2, 54, 59) but that so far have not been transformative in inducing breadth. In some cases, this may be because immunofocusing was not used (2, 59). As a result, successive shots may induce new strain-specific Abs, rather than building sufficiently on lineages initiated by preceding shots. In the other study (54), it is unclear why bNAbs did not develop. Possible explanations might be the repertoire limitations of rabbits for making V2 NAbs or insufficient shared memory T cell help between shots. The latter concern may be reduced for VLPs, as Gag should provide T cell help.

Radical strain changes are not a requirement for bNAb development in natural infection: bNAbs develop generally after a process of neutralization and escape over time with the infecting virus (24, 36, 53, 69). Superinfection with a new and diverse strain, is perhaps the closest natural infection scenario to our proposed SHPB regimens. Evidence suggests that superinfection doesn’t promote NAb breadth (130). However, an important difference in SHPB is that trimers are modified to immunofocus on NAbs, whereas superinfecting trimers are unlikely to share vulnerable sites. Depending on preliminary tests, to keep NAbs “on track” between shots, we may need to reduce or eliminate strain diversity. The variety of mutants of each strains will provide ample ways to reduce strain diversity in our regimens, as needed. Since early events are critical, it will be useful to define effective prime strategies and once cross-reactivity develops, diverse boosts may be powerful to promote further breadth.

In summary, we developed a diverse panel of membrane trimers with a range of V2 sensitivities. These trimers and the many variants of each will allow us to test a variety of vaccine concepts for immunofocusing V2 and FP NAbs. Ultimately, success in multiple polyclonal outbred models would provide strong support for clinical translation.

## Materials and Methods

### Plasmids

**i) HIV-1 Env plasmids.** Abbreviated names of Env strains are given first, with full names and GenBank references in parentheses, as follows: Q23 (Q23.17; AF004885.1), WITO (WITO.33, AY835451.1), c1080 (c1080.c3, JN944660.1), CM244 (CM244.ec1, JQ715397.1), T250 (also known as CRF250, T250.4, EU513189.1), 001428 (001428-2.42, EF117266.1), CE217 (CE703010217.B6-1, DQ056973.1), BB201 (BB201.B42, DQ187171.1), KER2018 (KER2018.11, AY736810.1), CNE58 (CNE58, HM215421.1), CAP45 (CAP45.G3, DQ435682.1), X2278 (X2278.c2.B6, FJ817366.1), JR-FL (JR-FL, AY669728.1), AC10 (AC10.29, AY835446.1), KNH1144 (KNH1144ec1, AF457066.1), sc422 (SC422661.8, AY835441.1), 6101 (6101, AY669708.1).

Full-length Env clones of the above strains, commonly used to make PVs for neutralization assays, were obtained from the NIH AIDS Reagent Repository, the Vaccine Research Center and The Scripps Research Institute. In many cases, these Env plasmids used expression plasmids such as pCI, pCDNA3.1, pCAGGS or pVRC8400. However, the modestly expressing plasmid pCR3.1 was used for Q23.17 and BB201.

**ii) Gag and Rev plasmids.** A plasmid expressing murine leukemia virus (MuLV) Gag (23). Whenever Env plasmids used native codons, we co-transfected pMV-Rev 0932 that expresses codon optimized HIV-1 Rev to maximize Env expression.

**iii) Glycosyltransferase plasmids** Glycosyltransferase plasmids pEE6.4_B4GalT1 (expressing β-1,4 galactosyltransferase and pEE14.4_ST6Gal1 (expressing β-galactoside α-2,6-sialyltransferase were co-transfected at a ratio of 1% and 2.5% total plasmid DNA, respectively.

**iv) MAb plasmids.** MAb plasmids were obtained from their producers and the NIH AIDS Reagent Repository. These included CH01/CH04, VRC38.01, PG9, PG16, and PGT145 directed to the V2 apex epitope; 39F and 14e directed to the V3 loop of gp120; VRC34.01 directed to the gp120-gp41 interface; and fusion peptide mAb vFP16.02 (8). UCAs of mAbs CH01/CH04 and VRC38.01 were described previously (13).

### VLP and gp120 monomer production

For VLP production, Env plasmids were co-transfected in Human Embryonic Kidney 293T or GnT1-293S cells using polyethyleneimine (PEI Max, Polysciences, Inc.), along with the MuLV Gag plasmid (23) and pMV-Rev 0932, as needed. 48 hours later, supernatants were collected, precleared, filtered, and pelleted at 50,000g in a Sorvall SS34 rotor. To remove residual medium, pellets were washed in 1ml of PBS, recentrifuged in a microcentrifuge at 15,000 rpm, and resuspended at 1,000x the original concentration in PBS. JR-FL gp120 monomer was produced and purified as described previously (77).

### Neutralization Assays

Assays were repeated at least twice to ensure consistency.

**i) NL-Luc assay.** Pseudoviruses (PV) were produced by co-transfecting 293T or 293S GnT1-cells with pNL4-3.Luc.R-E and an Env plasmid using PEI Max. Briefly, PV was incubated with graded dilutions of mAbs for 1 hour at 37°C, then added to CF2Th.CD4.CCR5 cells, plates were spinoculated, and incubated at 37°C (13). For wild-type (WT) PV, plates were incubated for 3 days, after which luciferase was measured. For SOS PV, following a 2-hour incubation, 5mM DTT was added for 15 minutes to activate infection. The mAb/virus mixture was replaced by fresh media, cultured for 3 days, and luciferase activity measured.

**ii) pQC-Fluc assay.** PV were produced by co-transfecting Env plasmids with pMLV GagPol and pQC-Fluc-dIRES (abbreviated as pQC-Fluc) (109). The resulting PV were used in neutralization assays with CF2Th.CD4.CCR5, as above.

**iii) Post-CD4 assay.** PV were mixed with sCD4 with or without V3 mAbs 14e or 39F. This mixture was then added to CF2.CCR5 cells, as described previously (102).

### Blue Native PAGE-Western Blot

VLPs were solubilized in 0.12% Triton X-100 in 1mM EDTA. An equal volume of 2x sample buffer (100mM morpholinepropanesulfonic acid (MOPS), 100mM Tris-HCl, pH 7.7, 40% glycerol, and 0.1% Coomassie blue) was added. Samples were spun to remove any debris and loaded onto a 4-12% Bis-Tris NuPAGE gel (Thermo Fisher) and separated for 3 hours at 4C at 100V. Proteins were then transferred to polyvinylidene difluoride (PVDF) membrane, de-stained, and blocked in 4% non-fat milk in PBST. Membranes were probed with a cocktail of mAbs 39F, 2F5, b12, 4E10, 14e, and PGT121, followed by alkaline phosphatase labeled anti-human Fc conjugate (Accurate Chemicals) and were developed using SigmaFast BCIP/NBT (Sigma).

### SDS-PAGE-Western Blot

VLPs were denatured by heating with 2-mercaptoethanol for 10 minutes at 90°C, then mixed with Laemmli buffer, then loaded onto 4-12% Bis-Tris NuPAGE gel (Invitrogen). To examine cleavage of oligomannose and hybrid glycans, endonuclease H (Endo H, New England Biolabs) was added to the samples after reduction and denaturation, followed by incubation for 1h at 37°C. Proteins were transferred onto PVDF membrane, de-stained, and blocked in 4% non-fat milk in PBST. Membranes were probed as for BN-PAGE blots above. Env band densities were quantified using Image Studio Lite (LI-COR).

### Reduction, alkylation and digestion of Env

JR-FL gp120 monomer was denatured for 1h in 50 mM Tris/HCl, pH 8.0 containing 6M urea and 5 mM dithiothreitol (DTT). Next, Env proteins were reduced and alkylated with 20mM iodoacetamide (IAA) for 1h in the dark, followed by a 1h incubation with 20mM DTT to eliminate residual IAA. Alkylated Env proteins were buffer exchanged into 50mM Tris/HCl, pH 8.0 using Vivaspin columns (3 kDa). Aliquots were digested separately overnight using trypsin and chymotrypsin (Mass Spectrometry Grade, Promega). The next day, the peptides were dried and extracted using C18 Zip-tip (Merck Millipore).

JR-FL VLPs were processed in the same way, except that were initially buffer exchanged into 50mM Tris HCl 0.1% Triton X-100 (w/w) to disperse lipids. To identify the glycome of the trimers that complexed with mAb CH01, VLPs were mixed with excess CH01 and incubated for 1h at 37°C. Screw cap spin columns were incubated with protein A–agarose for 10 minutes to allow for spin column resin equilibration before washing with gentle Ag-Ab binding buffer (Thermo Fisher Scientific). VLP-CH01 complexes were then applied to the spin columns and left to incubate for 30 minutes. Columns were washed twice with gentle Ag-Ab binding buffer prior to elution in 100-200 μL gentle Ag-Ab elution buffer (Thermo Fisher Scientific). Eluted VLP-CH01 mixtures were then buffer exchanged into 100μL 50mM Tris/HCl pH 8.0 for subsequent reduction and alkylation.

### Liquid chromatography-mass spectrometry (LC-MS) glycopeptide analysis

Peptides were dried again, re-suspended in 0.1% formic acid and analyzed by nanoLC-ESI MS with an Ultimate 3000 HPLC (Thermo Fisher Scientific) system coupled to an Orbitrap Eclipse mass spectrometer (Thermo Fisher Scientific) using stepped higher energy collision-induced dissociation (HCD) fragmentation. Peptides were separated using an EasySpray PepMap RSLC C18 column (75 µm × 75 cm). A trapping column (PepMap 100 C18 3μM 75μM x 2cm) was used in line with the LC prior to separation with the analytical column. LC conditions were as follows: 280 minute linear gradient consisting of 4-32% acetonitrile in 0.1% formic acid over 260 minutes, followed by 20 minutes of alternating 76% acetonitrile in 0.1% formic acid and 4% acetonitrile in 0.1% formic acid, to ensure all the sample elutes from the column. The flow rate was set to 300nL/min. The spray voltage was set to 2.7 kV and the temperature of the heated capillary was set to 40°C. The ion transfer tube temperature was set to 275°C. The scan range was 375−1500 m/z. Stepped HCD collision energy was set to 15%, 25% and 45% and the MS2 for each energy was combined. Precursor and fragment detection were performed with an Orbitrap at a resolution MS1=120,000, MS2=30,000. The AGC target for MS1 was set to standard and injection time set to auto which involves the system setting the two parameters to maximize sensitivity while maintaining cycle time.

### Site-specific glycan classification

Glycopeptide fragmentation data were extracted from the raw file using Byos (Version 3.5; Protein Metrics Inc.). Data were evaluated manually for each glycopeptide; the peptide was scored as true-positive when the correct b and y fragment ions were observed, along with oxonium ions corresponding to the glycan identified. The MS data was searched using the Protein Metrics 305 N-glycan library with sulfated glycans added manually. The relative amounts of each glycan at each site as well as the unoccupied proportion were determined by comparing the extracted chromatographic areas for different glycotypes with an identical peptide sequence. All charge states for a single glycopeptide were summed. The precursor mass tolerance was set at 4 ppm and 10 ppm for fragments. A 1% false discovery rate (FDR) was applied. The relative amounts of each glycan at each site as well as the unoccupied proportion were determined by comparing the extracted ion chromatographic areas for different glycopeptides with an identical peptide sequence. Glycans were categorized according to the composition detected.

HexNAc(2)Hex(10+) was defined as M9Glc, HexNAc(2)Hex(9−5) was classified as M9 to M3. Any of these structures containing a fucose were categorized as FM (fucosylated mannose). HexNAc(3)Hex(5−6)X was classified as Hybrid with HexNAc(3)Hex(5-6)Fuc(1)X classified as Fhybrid. Complex glycans were classified according to the number of HexNAc subunits and the presence or absence of fucosylation. As this fragmentation method does not provide linkage information, compositional isomers are grouped, so, for example, a triantennary glycan contains HexNAc(5) but so does a biantennary glycans with a bisect. Core glycans refer to truncated structures smaller than M3. M9glc-M4 were classified as oligomannose-type glycans. Glycans containing at least one sialic acid were categorized as NeuAc and at least one fucose residue in the “fucose” category.

Glycans were categorized into I.D.s ranging from 1 (M9Glc) to 19 (HexNAc(6+)(F)(x)). These values were multiplied by the percentage of the corresponding glycan divided by the total glycan percentage excluding unoccupied and core glycans to give a score that pertains to the most prevalent glycan at a given site. Arithmetic score changes were then calculated from the subtraction of these scores from one sample against others as specified.

### Construction of trimer model and cognate glycans

The model representation of the JR-FL SOS E168K+N189A trimer was constructed using SWISS-MODEL based on an existing structure of the 426c DS-SOSIP D3 trimer (pdb: 6MYY). Glycans were modelled on to this structure based on the most abundant glycoform identified from site-specific glycan analysis using WinCoot version 0.9.4.1 and PyMOL version 2.5.0. For sites which were not identified, a Man9GlcNAc2 glycan was modelled. Conditional color formatting was used to illustrate the predominant glycoforms of modeled glycans, as follows: green (high mannose), orange (hybrid) and magenta (complex).

### Phenotyping and calcium flux of CH01 UCA dKI-derived splenocytes

C57BL/6j WT or CH01UCA double KI (V_H_DJ_H_^+/+^ x VκJκ^+/+^) splenocytes were phenotyped with 0.5μg/mL of anti-B220 BV650, anti-CD19 APC-R700 (both from Becton Dickinson) and WITO-SOSIP-BV421 HIV Env tetramers, washed then stained with LIVE/DEAD® Fixable Near-IR Dead Cell Stain Kit (Thermo Fisher) for 30 min. To evaluate B-cell stimulation, splenocytes were stained with anti-B220 BV650 and anti-CD19 APC-R700 for 40 minutes. After washing with HBSS, pre-stained cells were loaded with Fluo-4 via by mixing with equal volumes of 2X Fluo-4 Direct™ loading solution (Fluo-4 Direct™ Calcium Assay Kit, Thermo Fisher). After a 30 min incubation at 37°C and then 30 mins at RT, cells were washed with HBSS and incubated with LIVE/DEAD Near-IR for 30 minutes. After another HBSS wash, cells were resuspended in calcium-containing HBSS and incubated at room temperature for 5 minutes before activation by anti-IgM F(ab′)2 (Southern Biotech) or VLPs. Fluo-4 MFI data for total B-cells (B220**^+^**CD19**^+^**) was acquired on a Beckman CytoFlex flow cytometer and analyzed using FloJo software.

### Negative-stain electron microscopy

A 4.8-µl drop of the sample was applied to a freshly glow-discharged carbon-coated copper grid for 10-15 s and removed using blotting paper. The grid was washed with several drops of buffer containing 10 mM HEPES, pH 7.0, and 150 mM NaCl, followed by negative staining with 0.7% uranyl formate. Staining quality and particle density were assessed using a Hitachi H-7650 transmission electron microscope. Representative images of VLPs were acquired with a Thermo Scientific Talos F200C transmission electron microscope operated at 200 kV and equipped with a Ceta CCD camera. The magnification was 57,000, corresponding to a pixel size of 0.25 nm.

#### Flow cytometry analysis of particles

Particle concentration, size, Env+ fraction and spike density were determined by single vesicle flow cytometry (113, 114), using a commercial kit (vFC^TM^ Assay kit, Cellarcus Biosciences, La Jolla, CA) and flow cytometer (CytoFlexS, Beckman Coulter, Indianapolis, IN). Briefly, samples were stained with the fluorogenic membrane stain vFRed^TM^ and anti-Env mAb PGT121, labeled with AlexaFluor647 (Thermo Fisher) for 1h at RT and analyzed using membrane fluorescence to trigger detection. Data were analyzed using FCS Express (De Novo Software), and included calibration using a vesicle size and fluorescence intensity standards. The analysis included a pre-stain dilution series to determine the optimal initial sample dilution and multiple positive and negative controls, per guidelines of the International Society for Extracellular Vesicles (ISEV) (129). A detailed description of vFC^TM^ methods and controls can be found in S3 Text. A MIFlowCyt Item Checklist and MIFlowCyt-EV, as required by the guidelines are provided in S1 Table.

## Acknowledgements

We thank John Mascola, Raiees Andrabi, Michael Farzan, Tommy Tong and Guillaume Stewart-Jones for suggestions and for providing reagents.

## Notes

### Competing Interest Statement

The authors have declared no competing interest.

